# *De Novo* Designed Peptide and Protein Hairpins Self-assemble into Sheets and Nanoparticles

**DOI:** 10.1101/2020.08.14.251462

**Authors:** Johanna M. Galloway, Harriet E. V. Bray, Deborah K. Shoemark, Lorna R. Hodgson, Jennifer Coombs, Judith M. Mantell, Ruth S. Rose, James F. Ross, Caroline Morris, Robert L. Harniman, Christopher W. Wood, Christopher Arthur, Paul Verkade, Derek N. Woolfson

**Affiliations:** School of Chemistry, University of Bristol, Cantock’s Close, Bristol, BS8 1TS, U.K.; School of Chemistry, University of Leeds, Leeds, LS2 9JT, U.K.; School of Biochemistry, University of Bristol, Medical Sciences Building, University Walk, Bristol, BS8 1TD, U.K.; BrisSynBio / Bristol Biodesign Institute, University of Bristol, Life Sciences Building, Tyndall Avenue, Bristol, BS8 1TQ, U.K.; Bristol Centre for Functional Nanomaterials, School of Physics, University of Bristol, HH Wills Physics Laboratory, Tyndall Avenue, Bristol, BS8 1TS, U.K.; School of Biological and Chemical Sciences, Fogg Building, Queen Mary University of London, Mile End Road, E1 4QD, U.K.; School of Chemistry, University of Glasgow, 0/1 125 Novar Drive, Glasgow, G12 9TA, U.K.; School of Biological Sciences, Roger Land Building, King’s Buildings, Edinburgh, EH9 3JQ, U.K.

**Keywords:** coiled coil, computational modelling, peptide design, protein design, self-assembly, synthetic biology

## Abstract

The design and assembly of peptide based materials has advanced considerably, leading to a variety of fibrous, sheet and nanoparticle structures. A remaining challenge is to account for and control different possible supramolecular outcomes accessible to the same or similar peptide building blocks. Here we present a *de novo* peptide system that forms nanoparticles or sheets depending on the strategic placement of a ‘disulfide pin’ between two elements of secondary structure that drive self-assembly. Specifically, we join homodimerizing and homotrimerizing *de novo* coiled-coil α-helices with a flexible linker to generate a series of linear peptides. The helices are pinned back-to-back, constraining them as hairpins by a disulfide bond placed either proximal or distal to the linker. Computational modeling and advanced microscopy show that the proximally pinned hairpins self-assemble into nanoparticles, whereas the distally pinned constructs form sheets. These peptides can be made synthetically or recombinantly to allow both chemical modifications and the introduction of whole protein cargoes as required.

## Introduction

Self-assembling proteins form the building blocks of life to control many, if not all, cellular process. Natural self-assembling proteins include: 1D actin filaments for cell structure and motility;^2^ 2D S-layers that form protective outer barriers in some bacteria;^3^ 3D capsids that protect viral DNA/RNA,^4^ and the shells of bacterial microcompartments, which are natural nanoreactors.^5^ These natural scaffolds can be engineered to display other proteins to create assemblies for biotechnology.^6, 7^ For example, enzyme pathways fused to viral capsid proteins self-assemble to create nanoreactors.^8^ Virus-like particles have also been engineered to display antigenic epitopes for use in vaccination,^9^ or adorned with targeting motifs to direct them to diseased cells.^10^ The toolbox of useful self-assembling protein structures has been expanded by mutating natural protein interfaces to induce controlled self-assembly and through *de novo* design.^11^ Such structures are usually computationally designed to form closely packed 2D arrays^12-18^, tubes^19-21^ or 3D icosahedral particles.^22-30^ However, these beautifully ordered and near crystalline assemblies may not be amenable to decoration with large cargos, or be permeable to small molecules due to their close packed nature. Also, these engineered or designed arrays can require thermal annealing to assemble,^15, 16^ which may reduce or even destroy the activity of any appended proteins. Alternatively, 1D fiber assemblies can form extensive 3D gels,^31, 32^ but controlling the localization of appended motifs and/or functions is limited because network formation is a stochastic process. Between these two extremes of close-packed order and 3D entangled networks, there is space for the development of room-temperature self-assembling biomolecular systems that can display functional cargos and be permeable to small molecules.

Previously, we have used *de novo* α-helical coiled-coil peptides to make self-assembled peptide cages (SAGEs).^33^ In SAGEs a homotrimeric coiled-coil peptide (CC-Tri3)^1^ is joined back-to-backwith a disulfide bond to one of two halves of a heterodimeric coiled-coil pair (CC-DiA and CC-DiB).^1^ This generates two complementary building-blocks or hubs, A & B.^33^ When mixed, these self-assemble into a hexagonal lattice that curves,^33^ and forms closed structures with the aid of defects.^34, 35^ SAGEs have been adapted for uptake by mammalian cells as potential drug delivery vehicles,^36^ and decorated with immunogenic peptide sequences to make a modular vaccine platform.^37^ The peptide cages are permeable to small molecules, so are ideal scaffolds for nanoreactors.^38^ The above examples all employ synthetically derived peptides and hubs. Natural proteins, such as enzymes and whole protein antigens, are too large to incorporate into SAGEs in this way. However, the system can be adapted to make peptide-protein fusions for recombinant production, leading to functionalized pSAGEs.^39^

Others have pioneered approaches to construct large coiled-coil based nanoparticles. For example, Burkhard and coworkers have linked a *de novo* homotrimeric and modified natural hompentameric coiled-coil sequences with a short GG linker and an inter-helix disulfide bridge proximal to the loop which self-assemble into polyhedral nanoparticles.^25, 28^ The group has been particularly successful at developing these as vaccine platforms for the presenation of antigenic peptides.^26-30^ In a different concept Marsh and colleagues combine *de novo* coiled-coil units and natural oligomeric proteins to render defined protein nanocages.^40-43^

Here we present self-assembling hairpin designs that form two distinct suporamolecular assemblies; namely, closed nanoparticles and extended sheets. These designs combine ideas from the SAGEs and Burkhard’s nanoparticles. Specifically, homodimeric (CC-Di) and homotrimeric (CC-Tri3) blocks^1^ are joined by a flexible linker, and pinned back-to-back with a disulfide bond (Figure 1). Computational modelling indicates that these should fold into stable hairpins that can be arrayed hexagonally Moreover, and distinctively, the models suggest that the position of the disulfide may influence the supramolecular assembly profoundly: a pin proximal to the loop leads to curved arrays, which could close to form particles, whereas a distal pin restricts curvature, potentially leading to extended sheets. These extremes are confirmed experimentally using a variety of advanced microscopy methods for hairpins made by peptide synthesis and when produced recombinantly. The validated designs offer *de novo* scaffolds with potential to display a range of functionalities for application in imaging, cell targeting, nanoreactors, drug delivery, and modular vaccines.

**Figure 1.**
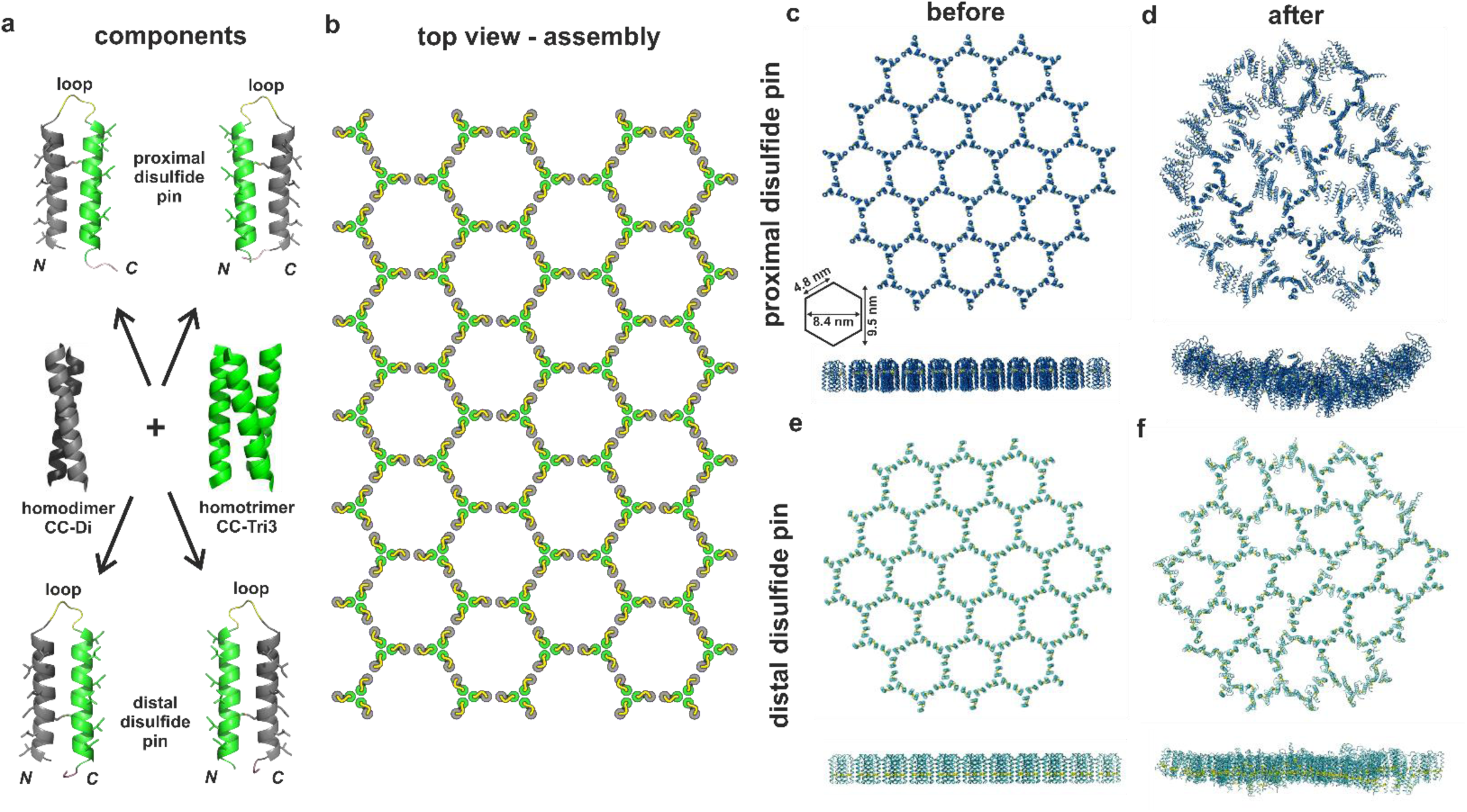
Rational design and assembly of helical hairpin peptide. (a) Schematic of hairpin design. The hairpins are constructed from two 3-heptad *de novo* coiled-coil peptides based on the homodimer CC-Di (grey, PDB code 4DZM) and the homotrimer CC-Tri3 (green, PDB code 4DZL). These are joined by a GSGSG ‘loop’ (yellow) and an interhelix disulfide bond ‘pin’ (black) between the polar helical facets. This leaves the coiled-coil forming facets exposed to engage in peptide-peptide interactions. The disulfide bond is placed either proximal or distal to the loop, which gives 4 possible hairpin configurations: with the dimer *N* terminal (left) or *C* terminal (right), with proximal (top) and distal (bottom) disulfide bonds. Full amino acid sequences for the designs are in Table 1. (b) Cartoon of the envisaged hexagonal network formed when the hairpins self-assemble *via* their coiled-coil interfaces. (c – f) Snapshots from molecular dynamics (MD) simulations of all-atom models of patches of assembled peptides, with dimensions of pre-simulated hexagon noted on (c). These were taken (c & e) before and (d & f) after 100 ns of MD under periodic boundary conditions using explicit TIP3P water, pH 7.0, 298 K, 0.15 M NaCl. Snapshots from assemblies of the (c & d) proximally pinned hairpins (Movie S1) and (e & f) distally pinned hairpins (Movie S2).

## Results and Discussion

### Design of α-Helical Hairpin Peptides

Our concept and design strategy is illustrated in Figure 1. Briefly, these used *de novo* α-helical coiled-coil peptides that form orthogonal assemblies with different oligomeric states as building blocks.^1^ Sequences for all-parallel homodimerizing (CC-Di) and homotrimerizing (CC-Tri3) peptides were linked linearly *via* a flexible GSGSG ‘loop’. Each coiled-coil sequence was modified to incorporate a cysteine at a normally exposed *f* position of the heptad repeat. The aim was for the two helical regions to be ‘pinned’ back-to-back in a hairpin arrangement. In turn, this should expose the hydrophobic *a* and *d* sites of each helical segment (opposite the *f* sites) to promote coiled-coil based self-assembly into hexagonal arrays. Two sites for the disulfide pins were explored in the full-length sequences: Cys15 plus Cys35 ‘proximal’ to the loop, and Cys8 plus Cys41 ‘distal’ from the loop, Figure 1a & Tables 1 & S1. The designs are named systematically, *e*.*g*. _pep_HP-DT_prox_ refers to the peptide (pep) hairpin (HP), with an *N*-terminal CC-Di and *C*-terminal CC-Tri3 (DT), and a proximal (prox) disulfide pin.

**Table 1.**
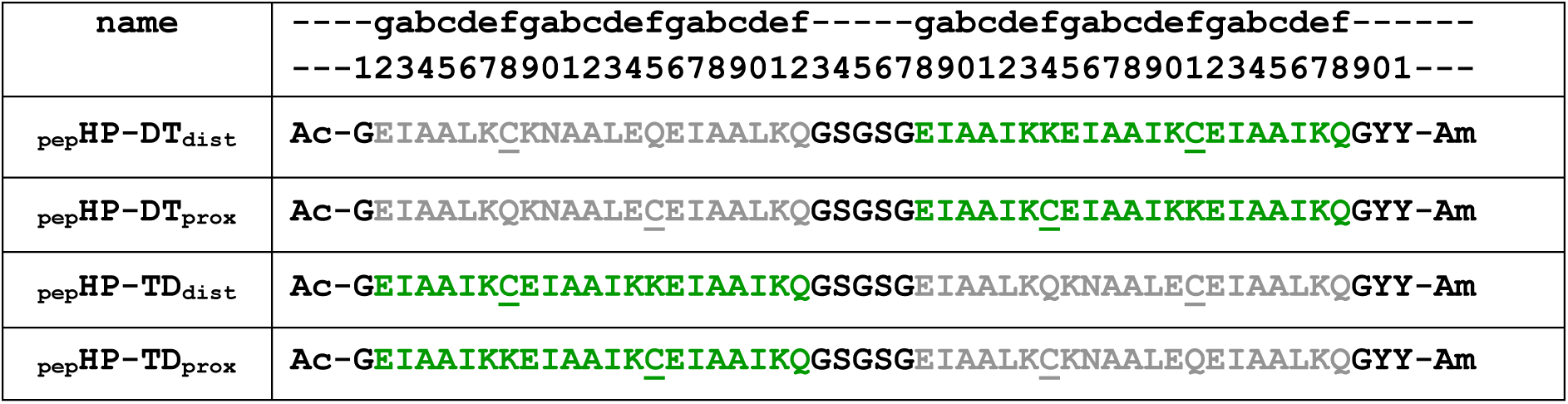
Peptide sequences synthesized using solid-phase peptide synthesis. Peptide name and corresponding amino acid sequence aligned with numbering of amino acid position (1 to 51) and coiled-coil register (*a* to *g*). Sequences based on a basis set of coiled-coils,^1^ with the sequence based on the homodimer (CC-Di) shown in grey, and the sequence based on the homotrimer (CC-Tri3) shown in green. Cysteine mutations introduced at f positions in order to form a disulfide bonded hairpin are underlined. Linkers (G, GSGSG) and chromophore tags (GYY) are shown in black.

### Modelling Indicates That Pin Position Influences Assembly Topology

To assess any differences in the placement of the disulfide pins, we constructed all-atom models for 19-hexagon patches of assembled arrays for the two different pin positions and subjected these to molecular dynamics (MD) simulations in water (Figure 1c-f, Movies S1 and S2). Intriguingly, the patch of the proximally pinned hairpins (_pep_HP-DT_prox_) curved in the 100 ns simulation. This resulted in the *N* and *C* termini of the peptides being presented on the convex face and the loops on the concave side. Projection of this curvature suggested that the arrays could close to form a nanoparticle 71 ± 11 nm in diameter, *i*.*e*. similar to the SAGEs (Figure S1a). In contrast, simulations of arrays of distally pinned hairpins (_pep_HP-DT_dist_),the 19-hexagon patches remained relatively flat throughout the trajectories, with no preferred direction or magnitude of curvature (Figure S1b). This suggested that these peptides might form flat extended sheets.

### Modelling Indicates That Pin Position Influences Assembly Topology

To assess any differences in the placement of the disulfide pins, we constructed all-atom models for 19-hexagon patches of assembled arrays for the two different pin positions and subjected these to molecular dynamics (MD) simulations in water (Figure 1c-f, Movies S1 and S2). Intriguingly, the patch of the proximally pinned hairpins (_pep_HP-DT_prox_) curved in the 100 ns simulation. This resulted in the *N* and *C* termini of the peptides being presented on the convex face and the loops on the concave side. Projection of this curvature suggested that the arrays could close to form a nanoparticle 71 ± 11 nm in diameter, *i*.*e*. similar to the SAGEs (Figure S1a). In contrast, simulations of arrays of distally pinned hairpins (_pep_HP-DT_dist_), the 19-hexagon patches remained relatively flat throughout the trajectories, with no preferred direction or magnitude of curvature (Figure S1b). This suggested that these peptides might form flat extended sheets.

These simulations gave the first indication of differences between the proximally and distally pinned systems. We rationalize this in terms of different flexibilities in the constructs. For the proximal pin, the helical termini have more freedom to explore space and force each other apart, resulting in a wedge-shaped building-block, narrow at the loop and wide at the termini. A distal pin reduces this freedom of movement and constrains the assembly to a flatter topology.

### Peptide Hairpins Assemble into Particles and Sheets as Predicted

All 4 peptide hairpins were synthesized, purified by HPLC, and confirmed as monomers by nanospray ionization mass spectrometry (Figures S2 & S3, Table S1). The peptides were oxidized with iodine to form the disulfide pin, and subject to Ellman’s test^44^ to confirm the disulfide bond formation. As judged by a change from clear to cloudy, all four peptides began to assemble as soon as they were hydrated in aqueous buffer. As a result, the samples scattered light and it was not possible to record reliable circular dichroism spectra to confirm α-helical assemblies.

Instead, we visualized the assemblies by negative-stain transmission electron microscopy (TEM) and cryogenic transmission electron microcopy (cryo-TEM), Figure 2. With either CC-Di or CC-Tri3 as the *N*-terminal block, the proximally pinned peptides _pep_HP-DT_prox_ and _pep_HP-TD_prox_ respectively, formed closed nanoparticle structures (Figure 2a-h). Both of the analogous peptides with the distal pin, _pep_HP-DT_dist_ and _pep_HP-TD_dist_, formed sheet-like structures, extending for >100 nm in the *xy* dimension (Figure 2i-p). Thus these initial experimental data support the MD simulations. Further, as there was no discernible difference in assemblies produced by the constructs with CC-Di or CC-Tri3 units placed first in the sequence, all further data presented had the CC-Di sequence at the *N*-terminus, *i*.*e*. HP-DT constructs.

**Figure 2.**
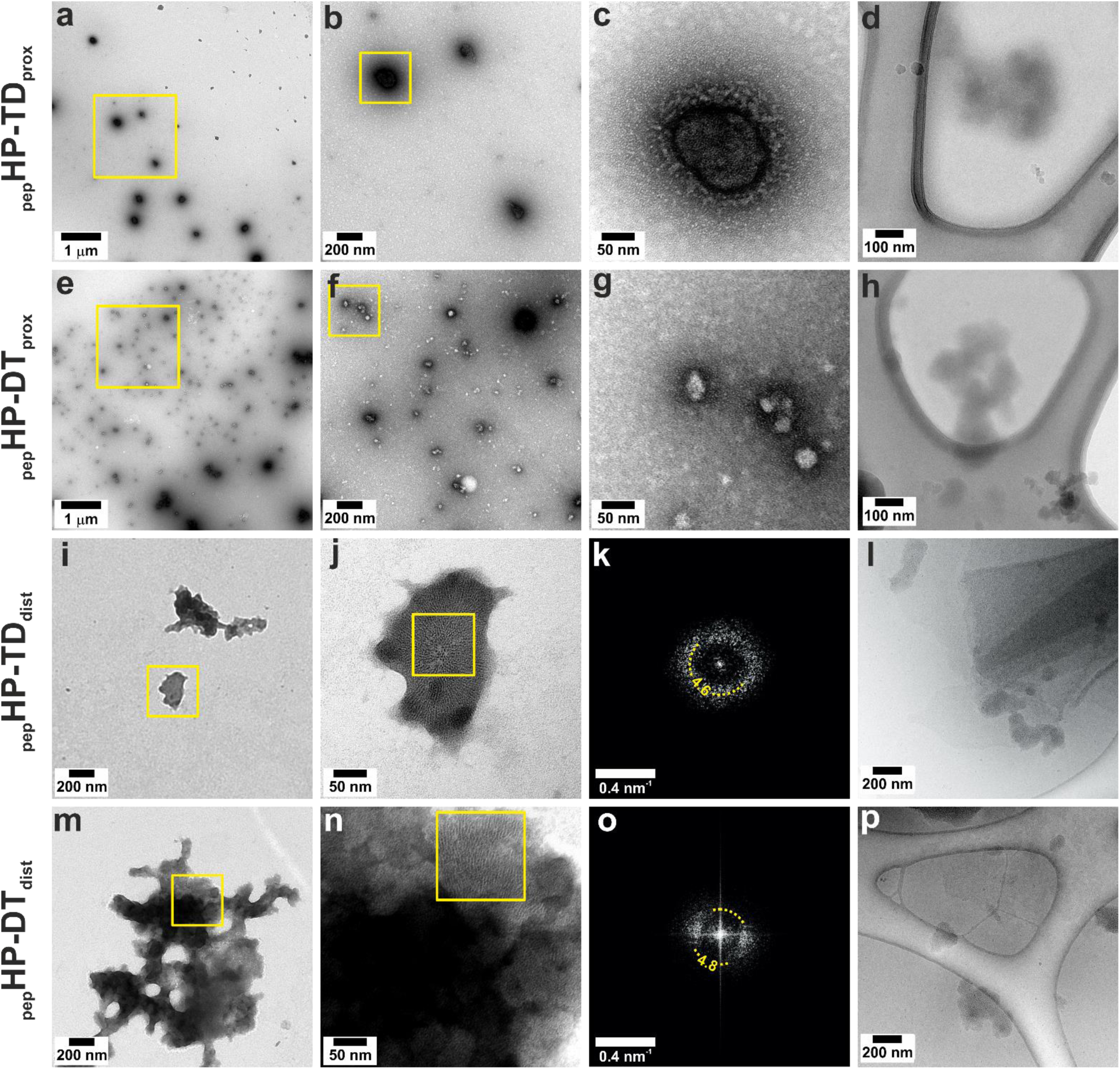
Negative stained TEM, FFTs, and cryo-TEM of peptide hairpin. Pin proximal to the loop: (a - d) _pep_HP-TD_prox_ and (e - h) _pep_HP-DT_prox_, formed closed structures. Pin distal to the loop: (i - l) _pep_HP-TD_dist_ and (m - p) _pep_HP-DT_dist_, formed flat structures. Negative stained samples at (a & e) low, (b, f, i & m) intermediate, and (c, g, j & n) high magnification. Fourier transforms (FFT) from distally pinned hairpins, area highlighted in (j) FFT shown (k) from _pep_HP-DT_dist_, and area highlighted in (n) FFT shown in (o) for _pep_HP-TD_dist_. (d, h, l & p) Cryo-TEM images. Hairpin peptide aliquots (100 µM) were assembled for 1 hour in HBS (25 mM HEPES, 25 mM NaCl, pH 7.2).

Next, we used atomic force microscopy (AFM) to probe the nature and dimensions of the assemblies in more detail, Figure 3. First, at 1 hour post assembly, the proximally pinned peptide _pep_HP-DT_prox_, formed nanoparticles with an average maximum height of 27 ± 22 nm and average diameter 101 ± 58 nm (n=5400), Figure 3a-c. The large ranges reflected a bimodal distribution of the aspect ratios, centered on ≈0.1 and ≈0.4. After 24 hours, the particles had approximately the same diameter (104 ± 79 nm), but only the thicker particles were apparent, with a height average of 39 ± 38 nm and aspect ratio 0.37, Figure 3d-f. Though the distributions were broad, these data are consistent with the particle sizes estimated from the modelling (71 ± 11 nm), and show that the particle diameters are stable over time. In addition, the experimental data indicated maturation of the particles between 1 and 24 hours. We cannot be sure what this is due to, but we posit that recruitment of peptides to the structures over time may result in organization into a multi-lamellar, stiffened particles, with the same diameter as seen in the earlier assembly.

**Figure 3.**
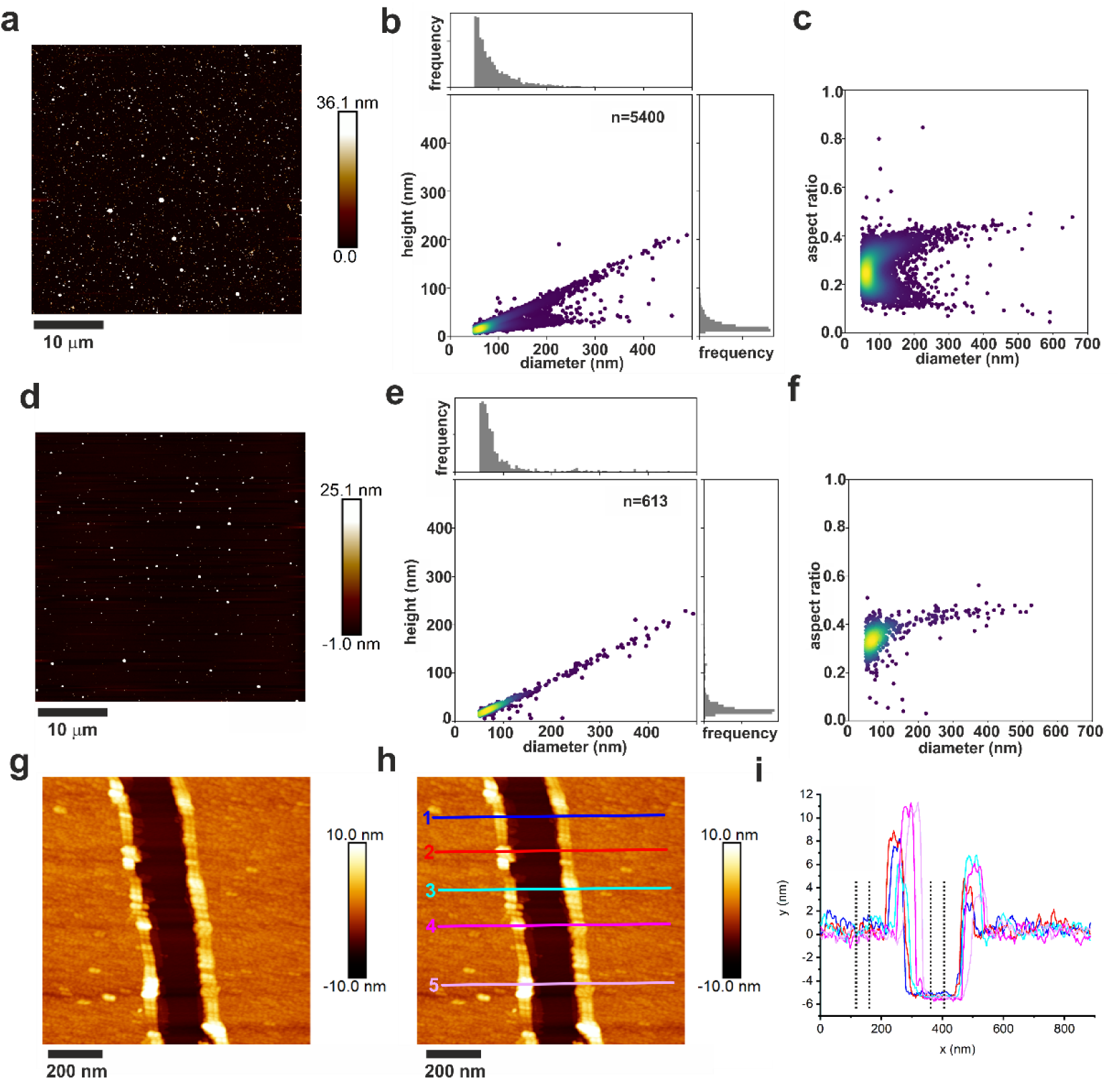
AFM grainsizing and height measurements of hairpin peptides. Proximally pinned _pep_HP-DT_prox_ (100 μM) were assembled in HBS (25 mM HEPES, 25 mM NaCl, pH 7.2).for (a - c) 1 hour and (d - f) 24 hours, then deposited onto mica. Images were recorded using peak force atomic force microscopy (PF-AFM, a & d). Particle height and diameter data were extracted using a particle analysis script (link available in Supplementary Methods) and plotted to show height versus diameter (b & e) and diameter versus aspect ratio (c & f). Distally pinned _pep_HP-DT_dist_ was assembled as above for 1 hour, deposited on mica and imaged using tapping mode AFM (TM-AFM, g). Height profiles were measured across a tear in Nanoscope analysis v1.5 (positions annotated on (h) and shown in (i)) where dotted lines indicate the area averaged to get the height difference between the mica substrate and the assembled peptide sheet, shown in Table S7.

AFM of the distally pinned peptide, _pep_HP-DT_dist_, revealed thin (5.6 ± 0.2 nm, Table S4) sheets (Figure 3g-i and S4). This is similar to the expected height of one hairpin from termini to the loop, which is ≈5 nm. This value, and the tight distribution of the experimental data support the hypothesis that these peptides form unilamellar sheets. It was not possible to discern the lattice clearly in AFM (Figure S5), which could be due to tip resolution, or flexibility and thermal motions in the assembly. However, fast Fourier transform (FFT) analysis of further TEM and cryo-TEM images (Figure 4b & d) revealed regular structures with spacings of 4.8 nm. This corresponds closely to the expected inter-vertex (inter-trimer) distances of the hexagonal lattice, which span 4 helices each just over ≈1 nm in diameter (Figure 1c).

**Figure 4.**
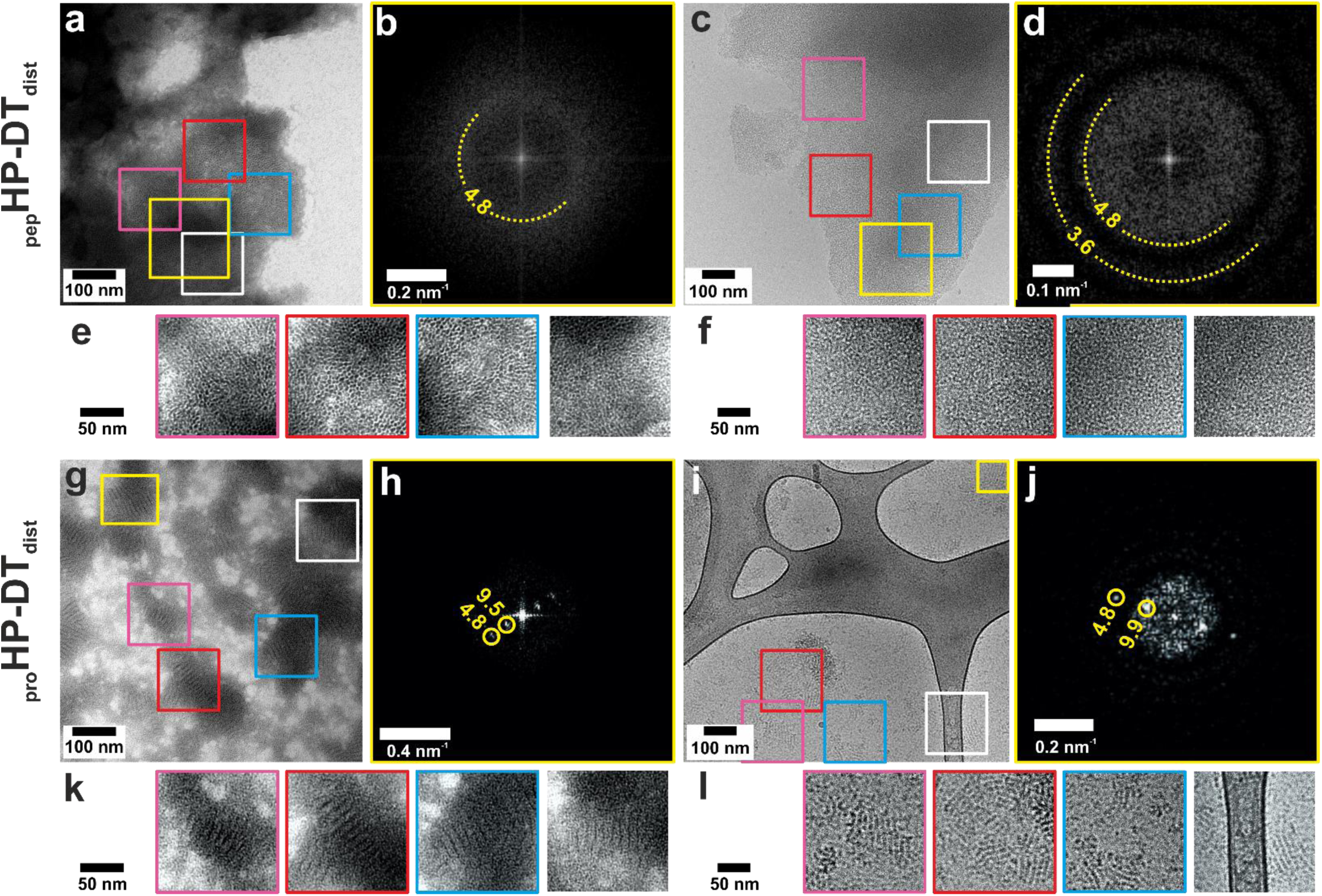
Negative stain TEM, cryo-TEM and FFTs showing peptide and protein hairpin sheet structures. Distally pinned hairpin peptide _pep_HP-DT_dist_ (a) negatively stained and (b) FFT of area indicated in yellow, and (c) cryo-TEM and (d) FFT of area indicated. Areas highlighted in (a) from negative stain are shown in (e), and highlighted in (c) from cryo-TEM are shown in (f), (enlarged 2x and brightness and contrast enhanced). Protein version _pro_HP-HT_dist_ (g) negatively stained and (h) FFT of area indicated, and (i) cryo-TEM and (j) FFT of area indicated. Areas exhibiting striped features of the assembled protein hairpins highlighted in (g) from negative stain are shown in (k), and highlighted in (i) from cryo-TEM are shown in (l) (enlarged 2x and brightness and contrast enhanced). Features in FFTs are labelled with their corresponding distance in real space in nm. Hairpin samples (100 µM) were hydrated in HBS (25 mM HEPES, 25 mM NaCl, pH 7.2) for 1 hour.

### Mixed Assemblies can be Made, but Pre-assembled structures do not Exchange Peptide Components

To test for peptide mixing during assembly and for exchange post assembly, we made two fluorescently labelled variants of _pep_HP-DT_dist_ and performed correlative light and electron microscopy (CLEM). We focused on this design as the sheets were much larger than the nanoparticles formed by the proximally pinned constructs, which made correlating the light and electron microscopy for individual sheets clearer than for the particles. The peptides had *N*-terminal carboxyfluorescein and 5(6)-carboxytetramethylrhodamine labels, Table S2. First, the labelled peptides were mixed when unfolded in 50% aqueous acetonitrile and freeze dried. When hydrated, these samples formed mixed assemblies as judged by the coincidence of the fluorescence from the two labels and electron density in CLEM, Figure S6 a-f. This demonstrates that differently decorated hairpin peptides can be combined into the same assembly. Second, green- and red-labelled hairpins were hydrated for 1 hour separately before mixing, incubated for 1 hour further, and then prepared for microscopy. In this case, the CLEM revealed distinct regions of green and red fluorescence, indicating that once assembled, the structures were stable and did not exchange peptide modules. Thus, despite their flexible construction, the hairpin peptides form robust and stable sheet assemblies from solution.

### Hairpin Redesign for Protein Expression

Next we sought to add functional proteins to the hairpin constructs. For this, we turned to the expression of synthetic genes in *Escherichia coli*. We designed genes for two constructs, _pro_HP-DT_prox_ and _pro_HP-DT_dist_, that harbored a 5’ multiple cloning site and a His-tag^45^ at the 3’ end (Figure S7, Tables S1 & S5); the prescript “pro” refers to the recombinantly expressed protein constructs. These were overexpressed in SHuffle ® T7 cells, which are engineered to allow disulfide bond formation in their cytoplasm, then purified and characterized by SDS-PAGE and confirmed as monomers by nanospray ionization mass spectrometry (Figure S8a-f). Ellman’s test^44^ confirmed that the molecules contained disulfide bonds. CLEM imaging of cell sections immunolabelled for the His-tag revealed protein dispersed within the cells rather than forming inclusion bodies (Figure S9).

Whereas the proximally pinned synthetic peptide hairpins formed spherical structures (Figure 2a-h), the recombinant variant, _pro_HP-DT_prox_ formed aggregates (Figure S10a pH 7.2). However, like the peptide variant (Figure 4a-f), the distally pinned protein hairpin, _pro_HP-DT_dist_ formed sheets (Figure 4g-l). Interestingly, these were smaller and had a clear ultrastructure, in the form of stripes. These stripes were apparent in both negative stain TEM (Figure 4e & k) and cryo-TEM (Figure 4f & l), and thus they are not a drying artefact and must reflect some underlying structure. FFTs of these revealed spots, with correspond with lines separated by ≈4.8 nm and ≈9.7 nm in real space in both cases (Figure 4h & j). As described above, this in consistent with the expected vertex-to-vertex spacing of ≈4.8 nm along the hexagon side, and of ≈9.5 nm across the 2 fold symmetry axis (Figure 1c).When compared to the peptide design, the protein has a flexible charge neutral region *N*-terminal to the hairpin sequence, and a *C*-terminal His-tag. It is possible that the small sheets are dimerizing through the His-tag,^46^ and forming slightly overlapping sheets, leading to Moiré fringes.^47^ Alternatively, these protein assemblies may form corrugated or stripy structures. Therefore we investigated how the protein patches behaved *in silico* and *in vitro* under different experimental conditions.

### pH Alters Protein Hairpin Assembly Structure

We constructed all-atom models for 19 hexagon patches of assembled arrays for the distally pinned and Histagged protein _pro_HP-DT_dist_. The pKa of the histidine side chain is near physiological pH.^48^ MD simulations we ran for the distally pinned protein hairpin _pro_HP-DT_dist_ with unprotonated and protonated His-tags as described above (Figure 5 and Movies S3 and S4). When unprotonated, the patch curved after 100 ns simulation (Figure 5b). The curvature was opposite to that seen for the proximally pinned peptide, with the loop on the convex side in unprotonated _pro_HP-DT_dist_ rather than on the concave side as was seen for _pep_HP-DT_prox_. The protonated constructs maintained a flat trajectory throughout the 100 ns simulation (Figure 5d), as was seen for the distally pinned peptide _pep_HP-DT_dist_.

**Figure 5.**
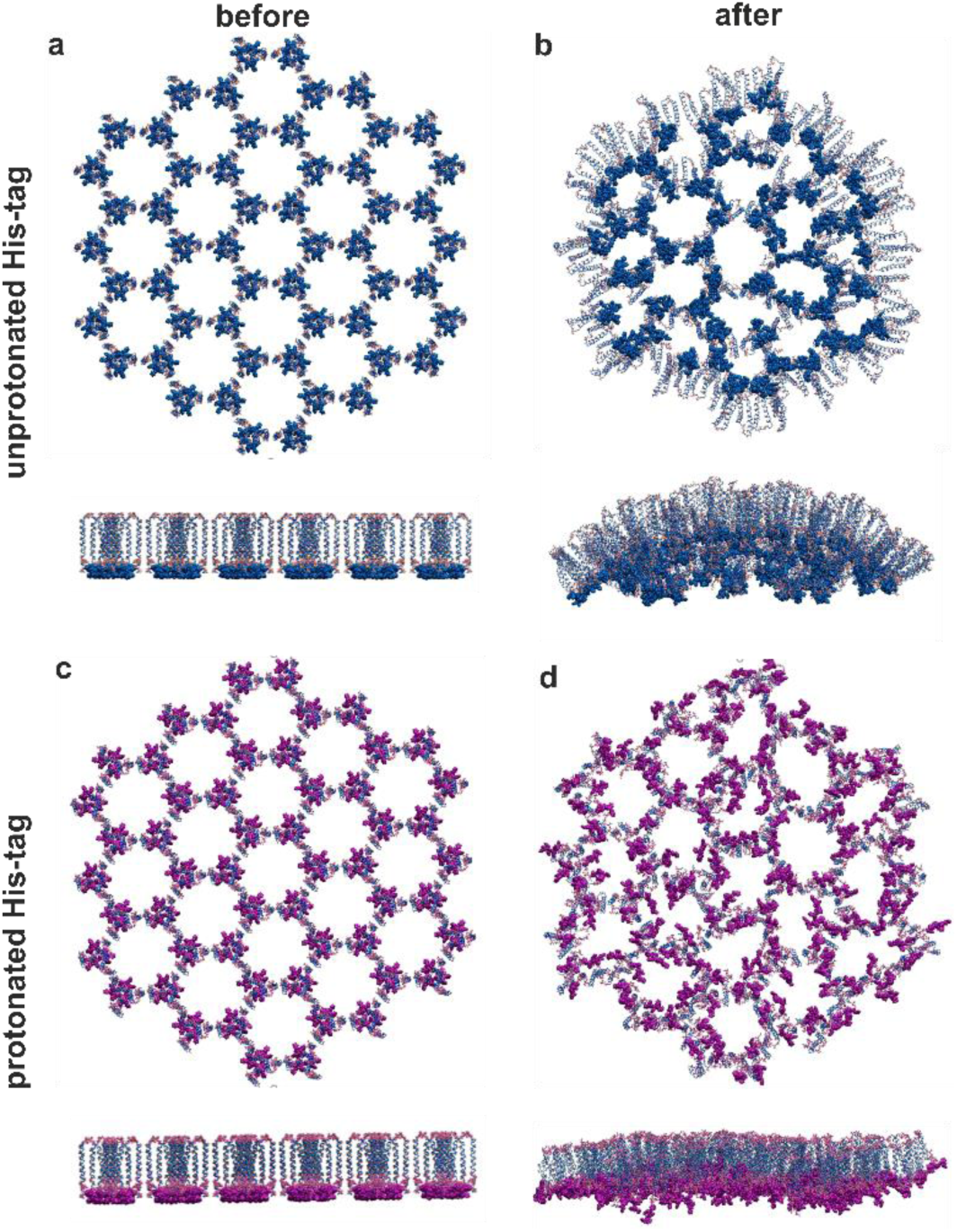
Snapshots from all atom MD simulations of 19 hexagon protein hairpin patches. Distally pinned protein hairpins (_pro_HP-DT_dist_) with (a & b) unprotonated (Movie S3) and (c & d) protonated (Movie S4) His-tags. Snapshots taken (a & c) before (blue, Movie S3) and (b & d) after (magenta, Movie S4) 100 nm of MD simulation under periodic boundary conditions using explicit TIP3P water, pH 7.0, 298 K, 0.15 M NaCl.

The assembly behavior of the protein hairpins was tested by hydration in HBS at pH values between 6.0 and 8.0 (Figure S10). The distally pinned protein formed extensive sheet like structures at low pH (<pH 6.5), and 3D aggregates at higher pH (8.0). Stripes appeared in the _pro_HP-DT_prox_ structures, but at slightly lower pH (6.5) when compared to the _pro_HP-DT_dist_ (7.2). Aggregates of _pro_HP-DT_prox_ were formed at low and high pH, indicating that the proximally pinned protein was not able to form stable structures. Modeling indicated that deprotonation of the His-tag induced the opposite sense of curvature to that caused by the proximal pin position. Thus, _pro_HP-DT_dist_ may be too flexible, and the opposite sense of curvature induced by protonation versus pin position destabilized the assembly leading to an aggregation based failure mechanism. The striped assemblies may be a transition state between the low pH “peptide-like” structure and the high pH “unprotonated curved” structures, probably induced by partial protonation of the His-tag. As its assembly was more stable, only _pro_HP-DT_dist_ was investigated further.

### Successful Self-assembly of Hairpins When Fused to Large Cargos

Fusions of the hairpin to the fluorescent protein sfGFP (∼28 kDa) (Figure S7, Table S1 & S5) were expressed in SHuffle ® T7 cells and purified as described above (Figure S8). The fluorescence in CLEM images of cell sections indicates that the sfGFP cargo folds in the cytoplasm (Figure S11). Again, we were not able to see any assembled structures within the cytoplasm. If the hairpins were able to self-assemble in cells, it may be that their open network structure is not electron dense enough to resolve in thin-section in TEM, so these constructs were also characterized *in vitro*. Ellman’s test^44^ confirmed that the molecules contained disulfide bonds after purification. As the nanospray mass spectrometry and SDS-PAGE also indicates _pro_GFP-HP-DT_dist_ forms a monomer (Figure S8g-i). When hydrated in HBS, we observed flat sheets formed by the cargo laden hairpins, (Figure S12). This demonstrates that distally pinned hairpins are amenable to being decorated with large proteins.

### Conclusions and Future Outlook

In summary, we have described the design, modelling, synthesis, assembly and characterization of two types of *de novo* peptide hairpins. In these, sequences for coiled-coil dimers and timers are joined through a flexible linker. The hairpins are then pinned, using disulfide bridges either proximal to or distal from the linker. The aim being to expose the hydrophobic seams of the coiled-coil segments and promote assembly of the hairpins into hexagonal arrays. The two pin positions have profoundly different effects on the topology of the supramolecular assemblies formed. The proximal pins lead to closed, spherical objects on the order of ≈100 nm in diameter, whereas the distal pin results in sheet-like assemblies consistent with a monolayer of folded, self-assembled hairpins. These structures are observed in a range of advanced microscopies, and the proposed mechanism of formation are supported by extended molecular dynamics simulations. Specifically, the proximal pin allows splaying of the helical termini, which in turn leads to curved arrays that can close, whereas the distal pins give a more tightly constrained hairpin structure, consistent with building blocks required to make flat sheets.

As noted above, others have developed similar self-assembling peptide and protein based nanoparticles^22-30^ or sheets.^12-18^ So what are the differences and advantages to our hairpin system?

First, by including two points (loop and pin) that define the relative positions and rotational freedom of the two coiled-coil components, we are able to control the topology of the self-assembled structures by design in a single system to render closed nanoparticles or extended sheets. This level of dual control is encouraging for future biomaterials design using relatively short linear peptides.

Second, although this is true for some other systems, the relatively short lengths of our hairpin designs and the minimal post-synthesis processing allows them to be made both synthetically and recombinantly. This has allowed us to decorate assemblies chemically, and through fusions to functional natural proteins; and to explore additional properties of the system. For example, experiments with fluorophore-labelled synthetic hairpin peptides show that co-assemblies can be made when starting with mixtures of unfolded peptide variants. However, once formed, there is no interchange between assembled structures. This approach could be used to add other functional moieties, *e*.*g*. for drug delivery and targeting diseased cells. Extending this to recombinantly produced hairpin-protein fusions, which express and purify well, could allow the production of nanoparticles or sheets with combinations of small molecule and protein cargoes.

Third, the fabric of the system is potentially modular, and we envisage that other *de novo* coiled-coil units could be swapped in to change the homomeric components used here.^49^ For instance, as we have demonstrated for the foregoing SAGE system,^33^ the homodimer sequence on the hairpins could be changed to a heterodimer pair.^49, 50^ This would afford an additional level of control on the assembly.

In all of these respects, the linear peptide hairpins that we describe provide *de novo* modules to add to the growing toolkit of components for the rational construction of biomaterials.^11, 24, 51, 52^ That said, how subtle differences in peptide design translate to relatively small changes in module structure and then to completely different assembly topologies are both alarming and encouraging. It is of concern because it highlights how careful the design process has to be in order to arrive at a targeted design, and that designs must be tested when appending functionality to any self-assembling building-block. However, it is also exciting, as it opens considerable possibilities for designing a wider range of biomaterial structures and functions. Whatever your stance, the rational and computational *de novo* design of biomaterials remains both challenging and full of potential.

## Materials and Methods

All materials and methods are listed in the supporting information available online.

## Additional Information

### Author Contributions

JMG, LRH, HEVB and DNW conceived the project and designed the experiments, JMG, HEVB, & JFR designed the peptide and protein sequences, JMG, HEVB & CM synthesized and purified the peptide sequences, JMG & RSR cloned the genes, and JMG expressed and purified the recombinant proteins. DKS set up, ran and analyzed the atomistic computational patch models of peptide assemblies. JMG, LHR, JC, JMM & PV recorded and analyzed TEM and cryo-TEM images. LHR & JMG processed cells for sectioning and staining, recorded fluorescent images and prepared CLEM alignments. HEVB, RLH, JMG, and CWW recorded and analyzed AFM images, and HEVB & CWW scripted the particle size analysis of AFM images. CA & JMG recorded and analyzed nanospray mass spectrometry data. JMG drafted the manuscript, which was written through contributions of all authors. All authors have given approval to the final version of the manuscript.

The authors declare no conflict of interest.

## Abbreviations

AFM: atomic force microscopy,
CC-Di: homodimeric coiled coil,
CC- Tri3: heterodimeric coiled coil from basis set sequences,^1^
cfl: 6-carboxyfluorescein,
CLEM: correlative light and electron microscopy,
cryo- TEM: cryogenic transmission electron microscopy,
dist: distal to the hairpin loop,
DNA: deoxyribonucleic acid,
*E. coli*: *Escherichia coli*,
FFT: fast Fourier transform,
FRET: Förster resonance energy transfer,
HEPES: 4-(2-hydroxyethyl)-1-piperazineethanesulfonic acid buffer,
HP: hairpin,
pep: peptide,
prot: protein,
prox: proximal to the hairpin loop,
SAGE: self-assembled peptide cage,
SDS-PAGE: sodium dodecylsulfide polyacrylamide gel electrophoresis,
sfGFP: super-folder green fluorescent protein,
tmr: TAMRA 5(6)-carboxytetramethylrhodamine,
TEM: transmission electron microscopy,
UA: uranyl acetate.

## Supporting information

This article contains supporting information online, which includes detailed experimental methods; DNA, protein and peptide sequences; cell strains used; protein and peptide characterization; supporting TEM, CLEM, AFM data; and movies and analysis of atomistic computational modelling of patches of assembled peptides and proteins. This material is available free of charge *via* the Internet. Data presented in this article will be made publically available on the Leeds Data Repository when the final version of the peer reviewed article is accepted..

## Acknowledgments

This research was supported by funding from a SLoLa (Strategic Longer and Larger) grant from the BBSRC “Development of supramolecular assemblies for enhancing cellular productivity and the synthesis of chemicals and biotherapeutics,” (BB/M002969/1), especially Martin Warren’s lab for supplying the pET3a and TBAD plasmids, and Dan Mulvihill for helpful discussion and insight, both based at the University of Kent.. We thank the EPSRC for awarding HECBiosym and an Archer Leadership Award for providing computer time on the U.K. supercomputer Archer. We thank the BBSRC/EPSRC funded Synthetic Biology Research Centre (BrisSynBio, BB/L01386X/1) for providing funding for researchers and Bluegem. We thank the EPSRC for funding the Chemical Imaging Facility (PF-AFM, EP/K035746/1), the mass spectrometry equipment (EP/K03927X/1), and the Woolfson Bioimaging Facility(fluorescence and electron microscopy BB/L014181/1) at the University of Bristol.

