## Supplementary material for "*De Novo* Designed Peptide and Protein Hairpins Self-assemble into Sheets and Nanoparticles": methods supplementary figures and tables

### EXPERIMENTAL

#### E1. Materials

Rink amide AM resin LL (100-200 mesh) for peptide synthesis, Fluorophores 5(6)-carboxyfluorescein and 5(6)-carboxytetramethylrhodamine were purchased from Novabiochem® (Merck, Watford, U.K), Cl-HOBt and Fmoc L-amino acids were from ACTG Bioproducts (Hessle, UK). DNA primers and inserts were ordered from GeneART® (Life Technologies Ltd., Thermo Fisher Scientific, Paisley, UK). Life Technologies Ltd. also supplied GeneJET plasmid miniprep kits, GeneJET gel extraction kits, protein standard ladders, His Tag monoclonal mouse / IgG2b antibody (MA1-21315), and Alexa fluor 488 labelled secondary polyclonal goat anti mouse IgG2b antibody (A32723). Superfolder GFP containing plasmid pBAD24-sfGFPx1 purchased from Addgene (#51558, Teddington, UK). DNA sequencing was performed by Eurofins (Wolverhampton, UK). Competent cells were purchased from Agilent Technologies (XL10-Gold®, Stockport, UK) and New England Biolabs (SHuffle® T7 Express, Hitchin, UK). IMAC Ni-NTA agarose resin was purchased from Qiagen (Hilden, DE). Anion exchange columns were purchased from GE Healthcare (Little Chalfont, UK). InstantBlue SDS-PAGE stain was purchased from Expedeon (CA, USA). Force microscopy probes: SCANASYST-AIR-HR (Bruker, CA, USA) were used for PeakForce (PF-AFM), and Scout-Beta (NuNano, Bristol, UK) for tapping mode (TM-AFM). All other reagents were purchased from Thermo Fisher Scientific (Loughborough, UK).

Aqueous buffers and solutions were made using ultrapure water (18.2 $M\Omega^{-cm}$ at 25 °C) and filtered using 0.22 μm sterile syringe filters purchased from Insight Biotechnology (London, UK). Formvar and carbon coated copper 200 mesh standard TEM grids, lacey carbon coated copper 200 mesh standard copper TEM grids and Lowicryl HM20 resin were purchased from EMS (Hatfield, PA, USA). Carbon-coated pioloform films on H6 finder grids were purchased from Agar Scientific (Stansted, UK).

#### E2. Molecular Dynamics Simulations

The coiled-coil helices were arranged as a 19-hexamer patch as previously described.[1] The proximal (Cys15 and Cys35) and distal (Cys8 and Cys41) hairpins were constructed by hand in InsightII (Accelrys) and each aligned to link one homodimer CC-Di sequence to one homotrimer CC-Tri3 sequence (movies S1 & S2), to which a *C*-terminal His-tag was appended if required (movies S3 & S4). In each case, residues comprising the core CC-Tri3 portion of CC-Di-hairpin-CC-Tri3 construct were aligned with each of the numbered hub peptides in the original 19-hexamer patch array using InsightII (Accelrys). Each aligned CC-Di-hairpin-CC-Tri3 peptide was then written out to provide coordinates to assemble as a new 19-hexamer patch. An in-house Fortran program was used to fix minor discrepancies in disulfide bond lengths. This was done for the patch with the hairpin pinned proximal to the loop (Cys15 and Cys35) and distal to the loop (Cys8 and Cys41) by moving the associated cysteine SG atoms towards each other, giving an S-S distance of 2.0 Å. This enabled the patch to retain the required 144 disulfides during the pdb2gmx process. Hydrogen atoms were added consistent with pH 7, and parameterized with the Amber-99SB9ildn force field. Each complex was surrounded by a box 4 nm larger than the assembled patch in each dimension, and filled with TIP3P water. Random water molecules were replaced by sodium and chloride ions to give a neutral overall charged box with an ionic strength of 0.15 M. Each box contained around 3 million atoms. Each patch was underwent 5000 steps of energy minimization prior to the molecular dynamics simulations.

The GROMACS-4.6.7 suite of software was used to set up and perform the molecular dynamics simulations. All simulations were performed as NPY ensembles at 298 K using periodic boundary conditions. Short range electrostatic and van der Waals' interactions were truncated at 1.4 nm, while long range electrostatics were treated with the particle-mesh Ewald's method, and a long range dispersion correction applied. Pressure was controlled by a Berendsen barostat, and temperature by the V-rescale thermostat. The simulations were integrated with a leap-frog algorithm over a 2 fs time-step, constraining bond vibrations with the P-LINCS method. Structures were saved every 0.1 ns for analysis and run over 100 ns. Simulation data were accumulated on the UK supercomputer Archer and the Bristol BrisSynBio supercomputer Bluegem. Molecular graphics manipulations and visualizations were performed using InsightII, VMD-1.9.1 and Chimera-1.10.2 and movies made using VMD. Curvature was measured at 19 points in the first 50 ns of the simulation (2-3 ns intervals) and used to extrapolate a particle size for the peptide hairpin simulations (Figure S1).

### E3. Peptide Synthesis and Purification

#### E3.1. Peptide Synthesis

Hairpins based on the basis set of α-helical coiled-coils[2] were synthesized using solid-phase peptide synthesis[3] (sequences shown in Table 1). A low loading rink amide resin support was used in a Liberty™ microwave peptide synthesizer. Fmoc protected amino acids (0.1 mM) were coupled (4.5 eq. 6-chloro-1-hydroxybenzotriazole (Cl-HOBt), 5 eq. *N,N'*-diisopropycarboydiimide (DIC) in 7 mL dimethylforamide (DMF) at 25 ºC for 2 minutes, then at 25 W microwave irradiation at 50 ºC for 5 minutes), washed (3x 7 mL DMF), deprotected (7 mL 20% (v/v) morpholine in DMF at 20 W and 75 ºC for 5 minutes), and washed (3x 7 mL DMF) before adding the next Fmoc protected amino acid.

#### E3.2. Peptide Capping

Resin mounted peptides were acetylated by incubating with 3 eq. acetic anhydride and 4.5 eq. *N,N*-diisopropylethylamine (DIPEA) in 10 mL DMF for 30 minutes at room temperature. To fluorescently label peptides, fluorophores were coupled to the *N*-terminal peptide in place of acetylation. Either 5(6)-carboxyfluorescein (cfl) or 5(6)-carboxytetramethylrhodamine (tmr) were coupled to the resin mounted peptide (5 eq. fluorophore, 4.5 eq. 1-[bis(dimethylamino)methylene]-1H-1,2,3-triazolo[4,5-b]pyridinium 3-oxid hexafluorophosphate (HATU), 5 eq. DIPEA, 10 mL DMF, 60 ºC, 2 hours). This was washed (5x 7 mL DMF), followed by incubation with 15 mL 20% (v/v) morpholine in DMF for 30 minutes at room temperature.

#### E3.3. Peptide Cleavage

Acetylated or fluorescently capped peptides were then washed (3x 5 mL DMF, 3x 5 mL dichloromethane (DCM)) and then dried. 9.5 mL trifluoroacetic acid (TFA), 250 µL water and 250 µL triisopropylsilane (TIPS) was mixed and then added to cleave the peptide from the resin, incubated on a mixer for 3 hours, then was separated by filtration. Excess solvent was evaporated using $N_2$, and the crude peptide precipitated using diethyl ether ($Et_2O$, 4 ºC). The peptide was pelleted by centrifugation (3000 rpm, 4 ºC, 10 minutes), the $Et_2O$ poured off and the pellet dissolved in 5-10 mL 50% (v/v) acetonitrile (MeCN) in water, and then freeze dried.

#### E3.4. Peptide Purification

Peptides were purified using reverse phase – high pressure liquid chromatography (RP-HPLC) using a Phenomenex C8 column (semi-micro, 5 µm, 100 Å, 10 mm x 250 mm). About 10 mg of crude peptide was dissolved in 3 mL containing 70% buffer A (0.1% (v/v) TFA in water) and 30% buffer B (0.1% (v/v) TFA in MeCN) and reduced by adding ≈10x excess tris(2-carboxyethyl)phosphine (TCEP, ≈8 mg) and incubating for 30 minutes. The reduced peptide was then injected onto the equilibrated column and a linear gradient between 30% and 70% buffer B was applied. Absorbance at 220 nm and 280 nm was used to monitor elution of the peptide from the column, indicating which peak to collect. The reduced peptide fractions were pooled and freeze dried. This was dissolved in 20% v/v aqueous acetic acid at ~1 mg $mL^{-1}$ (2-5 mL total volume) and 1-2 µL iodine solution added and rapidly mixed to oxidize the peptide and form the disulfide pin.[4] After 30-60 minutes, excess iodine was quenched by adding 10 µL aliquots of 100 mM sodium thiosulfate in water until the oxidized peptide solution turns from yellow to clear.[4] The solution was then freeze dried and dissolved in 2 mL (70% buffer A, 30% buffer B) and re-purified by RP-HPLC as above. Peptides were analyzed for purity by analytical RP-HPLC and for mass by nanospray ionization on a Waters SYNAPT G2S injected using an Advion TriVersa Nanomate® autoinjector. These data were fitted using Micromass MassLynx software maximum entropy analysis MaxEnt1 to 1 Da accuracy and compared to expected masses from pepcalc.com shown in Table S1.

#### E3.5 Quantitative Determination of Sulfhydryls

A quantitative Ellman's test[5] was performed to test that all of the peptide or protein in a purified and oxidized sample was disulfide pinned as expected. 4 mg of 5,5-dithiobis(2-nitrobenzoic acid) (DTNB) was dissolved in 1 mL 100 mM sodium phosphate buffer pH 8.0. The peptide or protein of interest was dissolved in 150 µL 100 mM sodium phosphate buffer (p 8.0) at a concentration of 50 µM. 5 µL of the DTNB solution was added and the mixture incubated for 5 minutes. A blank (buffer + DNTB solution) was also prepared. The absorbance of the reaction mixture was measured at λ = 410 nm, and the concentration of sulfhydryl groups calculated using the following Equation:[5]

$$[SH] = \frac{[A_{sample} - A_{blank}]}{13650}$$

where [SH] is the concentration of sulfhydryls (*i.e.* reduced cysteine side chains), and *A* is the absorbance at 410 nm. The oxidized peptides and proteins were checked against a fully reduced peptide hairpin at 50 µM concentration as a control.

### E.4. *In Vitro* Hairpin Assembly

Aliquots of between 1-10 nMol of peptide or protein were dissolved in 4-(2-hydroxyethyl)-1-piperazineethanesulfonic acid (HEPES) buffered saline (HBS, 25 mM HEPES, 25 mM NaCl, pH 7.2) to a final peptide/protein concentration of 100 µM (*e.g.* 10 nMol in 100 µL HBS = 100 µM solution) and were incubated to assemble for 1-24 hours at room temperature.

### E.5. Microscopy

#### E5.1. Atomic Force Microscopy (AFM)

5 µL of *in vitro* assembled hairpin was resuspended *via* pipette and dropped onto freshly cleaved mica. After ≈1 minute, excess liquid was wicked off with filter paper and the surface washed with 3x 100 µL water, which was wicked off and then gently dried using $N_2$ flow. PeakForce AFM (PF-AFM) imaging was done on a Multi-mode VIII microscope with a SCANASYST-AIR-HR cantilever (nominal tip radius 2 nm, spring constant 0.4 N $m^{-1}$, Brucker, CA, USA) on a fast scan head unit and a Nanoscope V controller. Tapping mode AFM (TM-AFM) was done using a Multi-mode using Scout-Beta cantilever (nominal tip radius 2 nm, spring constant 42 N $m^{-1}$, NuNano, Bristol, UK) with a Quadrexed Nanoscope III controller. Data was analyzed using Nanoscope Analysis 1.8 software (Brucker, CA, USA). Particle size analysis was performed using a script written by Dr. Chris Wood, available here: https://github.com/wells-wood-research/galloway-jg-hairpin-self-assembly-2020.

#### E5.2. *In Vitro* Assembled Samples for Transmission Electron Microscopy (TEM)

5 µL of assembled hairpin was resuspended *via* pipette and applied to a TEM grid (copper coated with 10 nm formvar and 1 nm carbon standard (FCF200-Cu), or Gilder finder (FCF200F2-Cu) grids, (EMS). After ≈1 minute, excess liquid was wicked away with filter paper, and grids washed with 10 µL water, which was then wicked off. 5 µL 1% (w/v) aqueous uranyl acetate was applied and incubated for ≈30 seconds before being wicked off. Finally, the samples were washed with 10 µL water, which was wicked off and the grids dried.

#### E5.3. *In Vivo* Assembled Samples for Transmission Electron Microscopy (TEM)

1 mL samples of cell culture were cooled to 4 ºC and pelleted (3000 xg, 4 ºC, 5 minutes), and the supernatant removed. 2 µL cell pellet was transferred to a 0.1 mm membrane carrier (Leica, Milton Keynes, UK) and frozen by high pressure freezing using an EMPACT2 + RTS system (Leica, Milton Keynes, UK). Vitrified carriers were freeze substituted with a solution of 0.2% (w/v) uranyl acetate and 5% (v/v) water in acetone in an automated AFS2 freeze substitution unit equipped with a FSP freeze substitution processor (Leica, Milton Keynes, UK), see Lee (2018).[6] Briefly, samples were held at -90 °C for 5 hours, then warmed at 5 °C $hour^{-1}$ to -45 °C, and held at -45 °C for 2 hours. Samples were washed 3x 30 minutes in etOH, then infiltrated with increasing dilutions (25%, 50%, 75%) of Lowicryl HM20 resin (EMS) in etOH for 3 hours each. Samples were then incubated in 100% Lowicryl HM20 resin for 16 hours, followed by 3x 2 hours incubation in fresh Lowicryl HM20 resin. The resin was polymerized using UV light (λ=360 nm) exposure throughout the following conditions: -45 °C for 16 hours, warmed at 5 °C $hour^{-1}$ to 0 °C (9 hours), then 14 hours further illumination at 0 °C.

Carriers were detached and the polymerized blocks trimmed prior to sectioning with a razor blade. 70 nm thick sections were cut using an EM UC6 microtome (Leica, Milton Keynes, UK) and a 45° diamond knife (DiATOME, PA, USA). Sections were collected onto carbon-coated pioloform films on H6 finder grids (Agar Scientific).

Immunolabelling of sections was done by first blocking grids by incubating on the top of a 100 µL drop of 1% (w/v) bovine serum albumen (BSA) in phosphate buffered saline (PBS, 140 mM NaCl, 2.7 mM KCl, 10.1 mM $Na_2HPO_4$, 1.8 mM $KH_2PO_4$ pH 7.4, 0.22 µm filter sterilized before use) for 2x 5 minutes. The grid was then incubated on a 10 µL drop of 1% BSA (w/v) in PBS containing 1:500 6x-His-tag monoclonal mouse / IgG2b antibody (MA1-21315, Life Technologies Ltd.). This was then washed by incubating on the top of 3x 100 µL drops of 0.1% BSA (w/v) in PBS for 5 minutes each, and then transferred to a 10 µL droplet drop containing DyLight 488 labelled secondary polyclonal goat anti mouse IgG antibody (35502, Life Technologies Ltd.) at 1 µg $mL^{-1}$ in 1% BSA (w/v) in PBS pH 7.4, and incubated for 1 hour. Grids were then washed by incubating on the top of a 100 µL drop of 0.1% BSA (w/v) in PBS for 5 minutes, then washed by incubating on the top of a 100 µL drop of water for a further 5 minutes.

#### E5.4. Correlative Light and Electron Microscopy (CLEM)[7]

Finder grids of fluorescent hairpin assemblies or cell sections were imaged in the bright-field and fluorescent channels on a Leica DMI4000B inverted epifluorescence microscope using a 63x oil objective lens with a numerical aperture 1.4. The same area of each sample was then imaged using TEM (see below). Fluorescence and TEM image data were aligned and overlaid using the TurboReg Registration plug-in in Fiji[8, 9] to create CLEM images.

#### E.5.5. Transmission Electron Microscopy (TEM)

After any fluorescence imaging, samples were imaged on a Tecnai 12 – FEI 120 kV BioTwin Spirit TEM (tungsten filament, accelerating voltage 120 keV), and images collected on a FEI Eagle 4k x 4k CCD camera. Images were processed using ImageJ[9, 10] including: adjusting contrast, annotating with scale bars, producing Fourier transforms (FFTs) and performing measurements.

#### E5.6. Cryo Transmission Electron Microscopy (Cryo-TEM)

Lacey carbon grids (LC325-Cu, EMS) were glow discharged for 30-60 seconds and mounted in a Leica EM GP automatic plunge freezer. 5 µL sample was applied to the grid within the humidity chamber (16 ºC, 90% humidity), blotted (1 second) and plunged into $N_2$ cooled liquid ethane. Grids were transferred to pucks and stored in liquid $N_2$. Frozen specimens were transferred to a Gatan CryoTransfer specimen holder and imaged using a Tecnai 20 – FEI 200 kV Twin Lens scanning transmission electron microscope (STEM) fitted with a $LaB_6$ filament. Low dose imaging (FEI low-dose module) minimized sample damage in the exposure area (spot size 3, 50,000x magnification) when using search mode (spot size 3, 2500x magnification) and focusing (spot size 3, 50,000x, 2-5 µm away from exposure area). Images were collected at ≈3 µm underfocus on a FEI Eagle 4k x 4k CCD camera. Higher resolution images were recorded on samples loaded into an FEI™ Talos™ Arctica equipped with a 200 kV X-FEG and a Gatan GIF Quantum LS energy filter on a Ceta 16 M CCD detector or Gatan K2 DED. Images were processed as for TEM (see above) as well as with the Gatan Microscopy Suite® (GMS) 3 software.

### E6. Protein Expression And Purification

#### E6.1. Plasmid Construction.

DNA sequences encoding $_{pro}$HP-HT$_{prox}$ and $_{pro}$HP-HT$_{dist}$ preceded with *Nco*I and *Kpn*I restriction sites and followed with a hexahistidine tag were optimized for production in *E. coil* and ordered from Gene-ART®. These were amplified by polymerase chain reaction (PCR, primers used in Table S3), extracted from agarose gels using a GeneJET gel extraction kit and cloned into the *Nde*I and *Spe*I sites of expression vectors pET3a (available from Merck, #69418) and TBAD (gifted by Warren Lab, School of Biosciences, University of Kent, UK) shown in Figure S6. A DNA insert encoding sfGFP[11] (pBAD24-sfGFPx1) was purchased from Addgene, and the fluorescent protein sequence was amplified and inserted 5' of the hairpin sequence using *Nco*I and *Kpn*I. Plasmids were amplified using XL10-Gold® competent cells (Agilent Technologies), and prepared for transformation into expression strains using a GeneJET plasmid miniprep kit (Life Sciences Ltd.). Cloning products were confirmed by DNA sequencing (Eurofins).

#### E6.2. Protein Expression

SHuffle® T7 Express competent cells (New England Biolabs), an *E. coli* strain engineered to facilitate disulfide bond formation in the cytoplasm, were transformed with the required plasmid. Transformants were plated onto LB agar (10 g L$^{-1}$ NaCl, 10 g L$^{-1}$ tryptone, 5 g L$^{-1}$ yeast extract, 16 g L$^{-1}$ agar) plus appropriate antibiotic (100 µg mL$^{-1}$ carbenicillin) to select for cells harboring the engineered plasmid. After incubation (37 ºC, 16 hours), a 10 mL starter culture of LB (10 g L$^{-1}$ NaCl, 10 g L$^{-1}$ tryptone, 5 g L$^{-1}$ yeast extract, pH 6.8) plus antibiotic (100 µg mL$^{-1}$ ampicillin) and 0.15% (w/v) D-glucose, and incubated (37 ºC, 16 hours) was inoculated with single colony. Protein was expressed using autoinduction media.[12] 400 mL LB was supplemented with antibiotic (100 µg mL$^{-1}$ ampicillin), and 8 mL 50x 52 (100 mL 50x autoinduction solution: 25 mL glycerol, 2.5 g D-glucose plus 10 g α-lactose (induction from pET3a vector, final concentration 0.05% glucose, 0.2% lactose) or 25 mL glycerol, 5.0 g D-glucose plus 5.0 g L-arabinose (induction from TBAD vector, final concentration 0.1% glucose, 0.1% arabinose)). Cultures were inoculated with 5 mL starter culture and incubated (30 ºC, 200 rpm) for up to 24 hours. 1 mL samples of cultured cells were collected for preparation for imaging (see *in vivo* assembled samples above). Cells for protein purification were pelleted (10,000 xg, 4 ºC, 10 minutes), the supernatant discarded and the pellets frozen at -20 ºC.

#### E6.3. Protein Purification

Frozen cell pellets were thawed and resuspended in 20 mL sonication buffer (TBS: 50 mM Tris, 150 mM NaCl, pH 8.0; plus: 100 µM ethylenediaminetetracetic acid (EDTA), 0.5 mM phenylmethane sulfonyl fluoride (PMSF), and 1% (v/v) triton x100) and sonicated on ice (1 second burst, 1 second rest for 15 minutes). Lysates were cleared (29,000 xg, 4 ºC, 10 minutes), and the protein purified from the supernatant by immobilized metal ion affinity chromatography (IMAC).[13] The supernatant was applied to a Ni-NTA agarose resin (Qiagen) to immobilize the protein of interest *via* the His-tag. 20 mL wash buffer (50 mM Tris, 150 mM NaCl, 20 mM imidazole, pH 8.0) was applied to remove loosely bound proteins, followed by 20 mL elution buffer (50 mM Tris, 150 mM NaCl, 300 mM imidazole, pH 8.0) to release the protein of interest, which was collected in 2 mL fractions.

Fractions containing the eluted protein were analyzed for purity using SDS-PAGE (see below), pooled, and further purified by anion exchange on an ÄKTAprime plus (GE Healthcare). Smaller proteins were applied to a HiTrap DEAE FF low bind anion exchange column, and larger proteins to HiTrap Q FF high binding column (both 5 mL, GE healthcare). A gradient between a low salt buffer (50 mM Tris, 25 mM NaCl) and a high salt buffer (50 mM Tris, 500 mM NaCl) was applied at 2 mL per minute over 30 minutes. Absorbance at $\lambda$ = 280 nm was monitored as a proxy for protein concentration, and eluted protein was collected in 2 mL fractions. Fractions were analyzed by SDS-PAGE (Figure S8) for purity and protein mass. Suitable fractions were pooled, buffer exchanged into 50 mM ammonium bicarbonate pH 7.9, concentrated and freeze dried in 1, 5, & 10 nM aliquots. Smaller proteins (<10 kDa) could also be purified by RP-HPLC as described above for peptide purification where required.

#### E6.4. Protein Analysis

Protein samples from whole cells, soluble mixtures and Ni-NTA, anion exchange and RP-HPLC fractions were analyzed using sodium dodecyl sulfate – polyacrylamide gel electrophoresis (SDS-PAGE) under denaturing conditions. Smaller target proteins (<10 kDa) were run on tricine gels[14] with a low molecular weight ladder (3.4-100 kDa range, Life Sciences Ltd.) whereas larger target proteins were run on gels with glycine running buffer[15] with a standard PAGE ruler (10-180 kDa range, Life Sciences Ltd.). For gel compositions, see Table S6. Protein samples were mixed with 4x loading buffer, heated at 95 ºC for 5 minutes, then centrifuged at 17,000*xg* for 5 minutes. For tricine gels, separate cathode and anode buffers were used, whereas a single running buffer was used for glycine gels (see Table S7). 5-15 µL sample was loaded into each well, and 180 V applied for 40-50 minutes until the dye front reached the bottom of the gel. Gels were either stained with InstantBlue (Expedeon) and destained with water, or stained with Coomassie blue and revealed with destain (see Table S7).

### SUPPLEMENTARY FIGURES AND TABLES

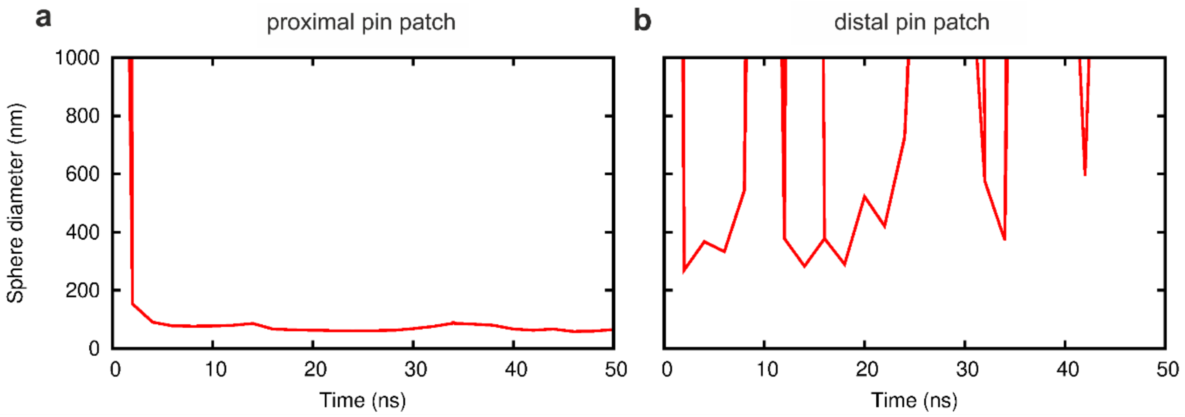

**Figure S1. Plots of curvature against time extracted from simulation trajectories.** Curvature is used to extrapolate a particle diameter from the modelled patch for (a) the proximally pinned hairpin $_{pep}$HP-DT$_{prox}$ and (b) the distally pinned hairpin $_{pep}$HP-DT$_{dist}$. (a) Shows that the curvature of the simulated proximally pinned hairpin patch settles to a value consistent with forming a sphere 71.4±11.4 nm in diameter (average and standard deviation of values measured between 5 ns and 50 ns). (b) Shows that the distally pinned hairpin patch was not able to curve, but instead the expected particle fluctuates between high values (100s of nm) and infinity, indicating a flat rather than closed structure would be formed by these constructs.

**Table S1. Peptide and protein name, sequences, mass and pI**. Dimer (CC-Di) shown in grey, and trimer (CC-Tri3) shown in green. Cysteine at *f* positions are underlined. Peptide caps: Ac = acetylated, Am = amidated; cfl = 5(6)-carboxyfluorescein; tmr = (5-carboxtetramethylrhodamine). Fluorescent protein sequence GFP in orange. Predicted mass of oxidized sequence, mass measured by nanospray and pI calculated using pepcalc.com (Innovagen, Lund, SE). Upper values for proteins are for the full length sequence, lower value is missing the *N*-terminal methionine, a common occurrence for proteins where serine (S) occurs after the initial methionine (M).[16]

| name | sequence | predict mass Da | measure mass Da | pI |
|---|---|---|---|---|
| $_{pep}$HP-DT$_{dist}$ | **Ac-GEIAALKCKNAALEQEIAALKQGSGS-**<br>**GEIAAIKKEIAAIKCEIAAIKQGYY-Am** | 5317.2 | 5317 | 8.11 |
| $_{pep}$HP-DT$_{prox}$ | **Ac-GEIAALKQKNAALECEIAALKQGSGS-**<br>**GEIAAIKCEIAAIKKEIAAIKQGYY-Am** | 5317.2 | 5317 | 8.11 |
| $_{pep}$HP-TD$_{dist}$ | **Ac-GEIAAIKCEIAAIKKEIAAIKQGSGS-**<br>**GEIAALKQKNAALECEIAALKQGYY-Am** | 5317.2 | 5316 | 8.11 |
| $_{pep}$HP-TD$_{prox}$ | **Ac-GEIAAIKKEIAAIKCEIAAIKQGSGS-**<br>**GEIAALKCKNAALEQEIAALKQGYY-Am** | 5317.2 | 5316 | 8.11 |
| $_{pep}$cfl-HP-DT$_{dist}$ | **cfl-GEIAALKCKNAALEQEIAALKQGSGS-**<br>**GEIAAIKKEIAAIKCEIAAIKQGYY-Am** | 5633.4 | 5633 | 8.11 |
| $_{pep}$tmr-HP-DT$_{dist}$ | **tmr-GEIAALKCKNAALEQEIAALKQGSGS-**<br>**GEIAAIKKEIAAIKCEIAAIKQGYY-Am** | 5687.6 | 5687 | 8.11 |
| $_{pro}$HP-DT$_{dist}$ | **MSGSMGSSGT-**<br>**GEIAALKCKNAALEQEIAALKQGSGS-**<br>**GEIAAIKKEIAAIKCEIAAIKQGYYGSGHHHHHH** | 7183.1<br>7051.9 | 7055 | 8.08 |
| $_{pro}$HP-DT$_{prox}$ | **MSGSMGSSGT-**<br>**GEIAALKQKNAALECEIAALKQGSGS-**<br>**GEIAAIKCEIAAIKKEIAAIKQGYYGSGHHHHHH** | 7183.1<br>7051.9 | 7052 | 8.08 |
| $_{pro}$GFP-HP-DT$_{dist}$ | **MSGSMGMSKGEELFTGVVPILVELD-**<br>**GDVNGHKFSVRGEGEGDATNGKLTLKFICTT-**<br>**GKLPVPWPT-**<br>**LVTTLTYGVQCFSRYPDHMKRHDFFKSAMPEGYVQERT**<br>**ISFKDDGTYKTRAEVKFEGDTLVNRIELKGID-**<br>**FKEDGNILGHKLEYNFNSHNVYIT-**<br>**ADKQKNGIKANFKIRHN-**<br>**VEDGSVQLADHYQQNTPIGDGPVLL-**<br>**PDNHYLSTQSVLSKDPNEKRDHMVLLEFVTAAGITH-**<br>**GMDELYKGT-**<br>**GEIAALKCKNAALEQEIAALKQGSGS-** | 33773.8<br>33640.6 | 33647 | 6.39 |

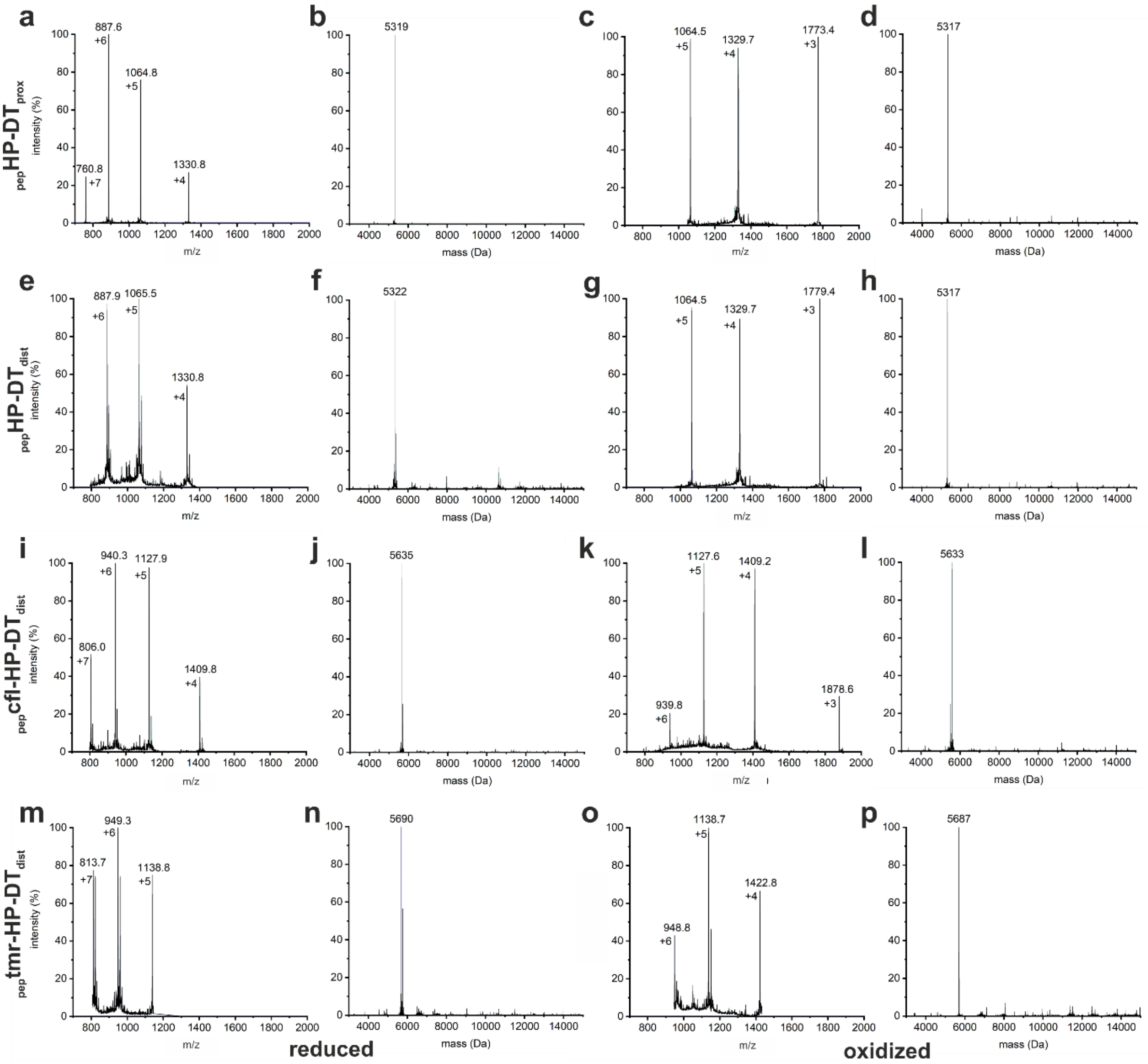

**Figure S2**. **Example plots of nanospray ionization mass spectrometry of peptides** synthesized for this study. Peptides after reduction with TCEP and first purification as the (a, e, i & m) raw data and (b, f, j & n) processed data, and after oxidation with iodine and the second RP-HPLC purification as the (c, g, k & o) raw data and (d, h, l & p) processed data. Peptides shown are (a - d) $_{pep}$HP-DT$_{prox}$, (e - h) $_{pep}$HP-DT$_{dist}$, (i - l) $_{pep}$cfl-HP-DT$_{dist}$, and (m - p) $_{pep}$tmr-HP-DT$_{dist}$. Reduced peptides show mass of peptide as predicted by pepcalc.com, and oxidized peptides show mass of 2 Da less than this as two hydrogens on cysteine are lost as the disulfide forms to create the pin. Data for all reduced peptides and purified proteins are shown in Table 1 & S1.

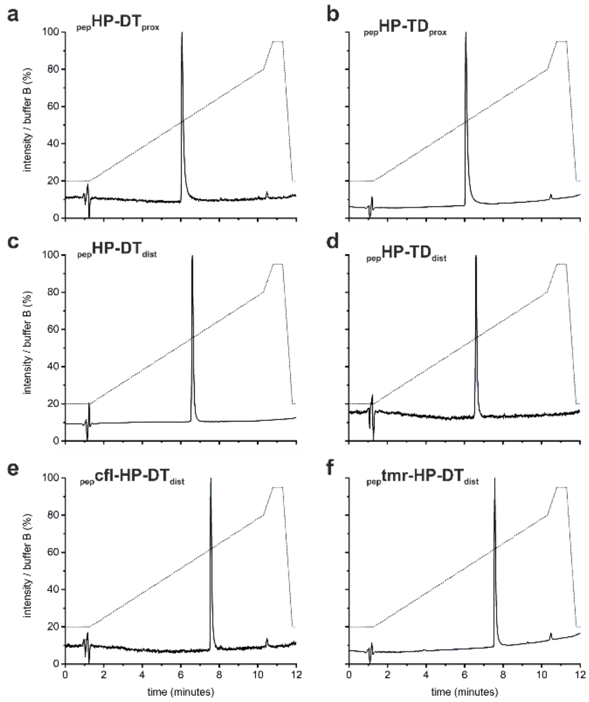

**Figure S3. Analytical RP-HPLC of peptides hairpins** after initial oxidation to form the hairpin, and subsequent final purification by RP-HPLC. Absorbance at a wavelength of λ = 220 nm is shown as a solid trace, and gradient applied as a percentage of buffer B is shown as a dotted line. (a) $_{pep}$HP-DT$_{prox}$, (b) $_{pep}$HP-TD$_{prox}$, (c) $_{pep}$HP-DT$_{dist}$, (d) $_{pep}$HP-TD$_{dist}$, (e) $_{pep}$cfl-HP-DT$_{dist}$, and (e) $_{pep}$tmr-HP-DT$_{dist}$.

**Table S2**. **Height of hairpin peptide sheet measured from AFM** tapping mode image shown in Figure 3. Measurements are taken as the difference between the mica substrate and the peptide sheet away from the edge where the sheet appears to have curled up.

| **profile** | **height (nm)** |
|---|---|
| 1 | 5.94 |
| 2 | 5.70 |
| 3 | 5.45 |
| 4 | 5.54 |
| 5 | 5.41 |
| **average** | **5.61 ± 0.22** |

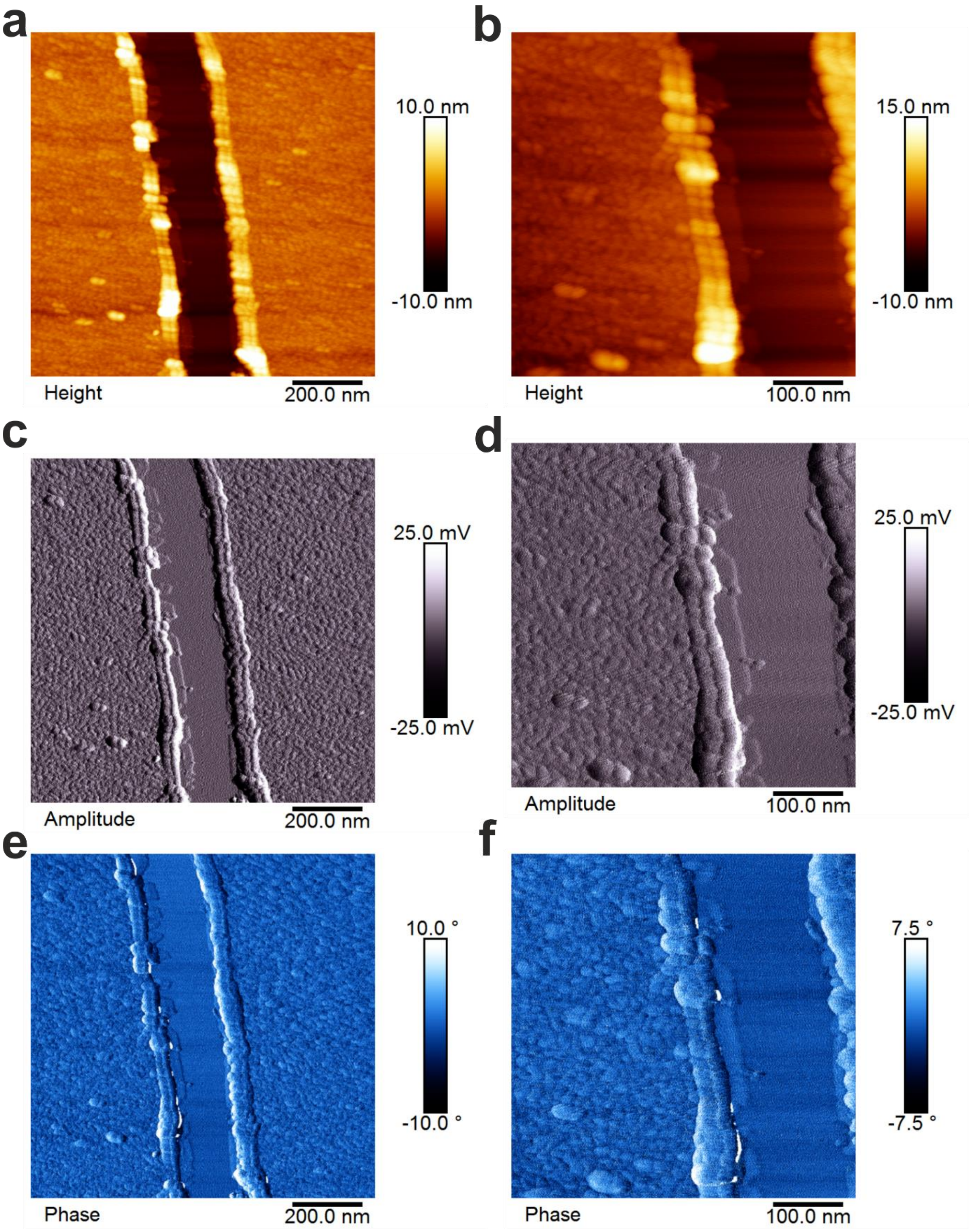

**Figure S4. Tapping mode atomic force microscopy (TM-AFM) images of $_{pep}$HP-DT$_{dist}$ peptide sheet**. An area was selected that shows a tear in the self-assembled peptide sheet, with the mica substrate appearing smooth (central area) when compared to the peptide sheet areas (left and right sides of image) at 2 magnifications. (a & d) show the topography, with the height of the sheet at about 10 nm. The same feature can be seen in the amplitude (b & e) and phase shift (c & f) plots.

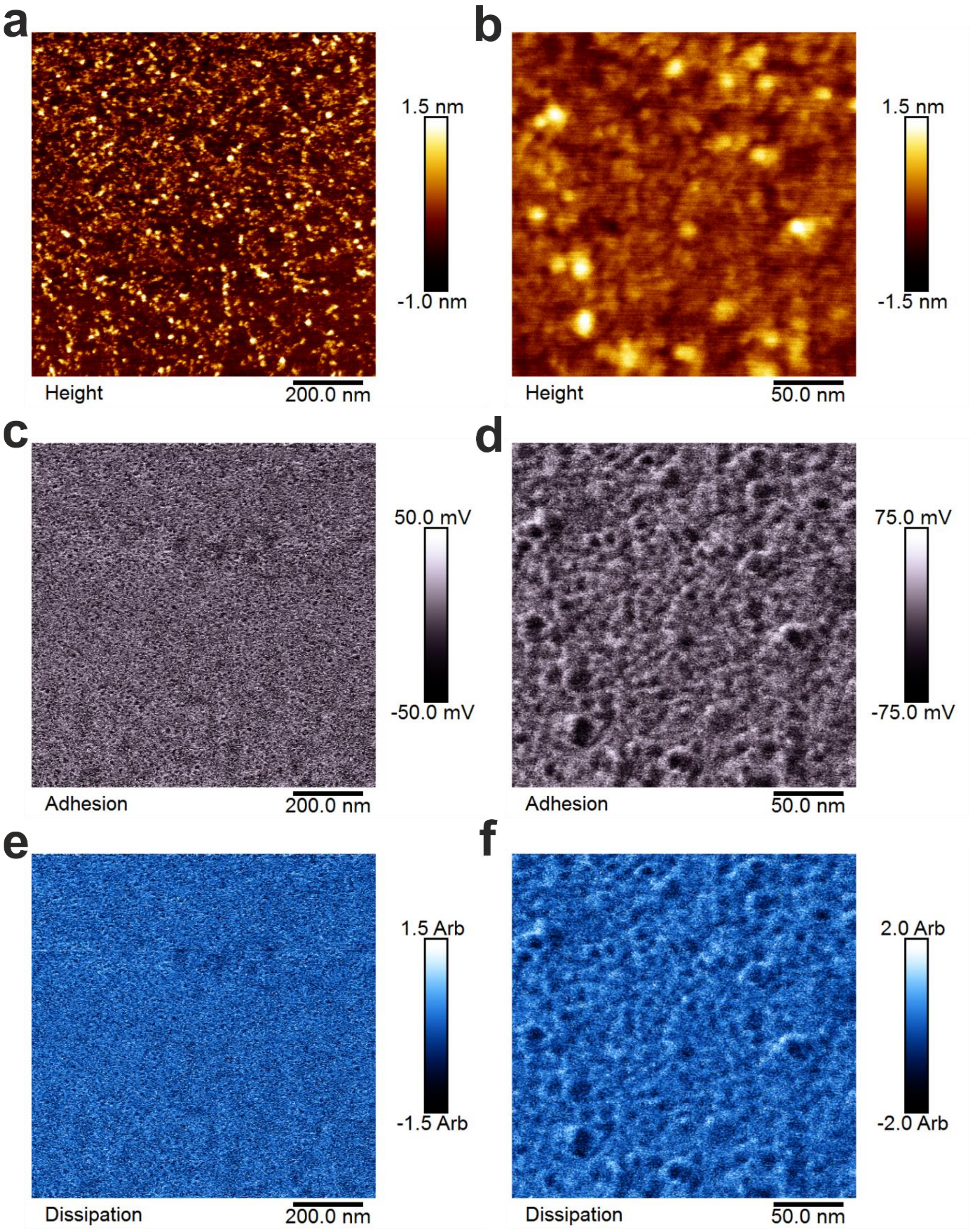

**Figure S5. Peak force atomic force microscopy (PF-AFM) images of $_{pep}$HP-DT$_{dist}$ peptide sheet.** An area was selected was in the middle of a self-asssembled sheet. (a & d) Show the topography, with the variability in height of the sheet at about 2 nm (including the spots, which are likely to be salt crystals). The same features can be seen in the amplitude (b & e) and phase shift (c & f) plots. At the highest magnification (c-f) some latticework features can be seen. However, the image resolution and/or order in the pattern is not high enough to resolve features *e.g.* as a Fourier transform.

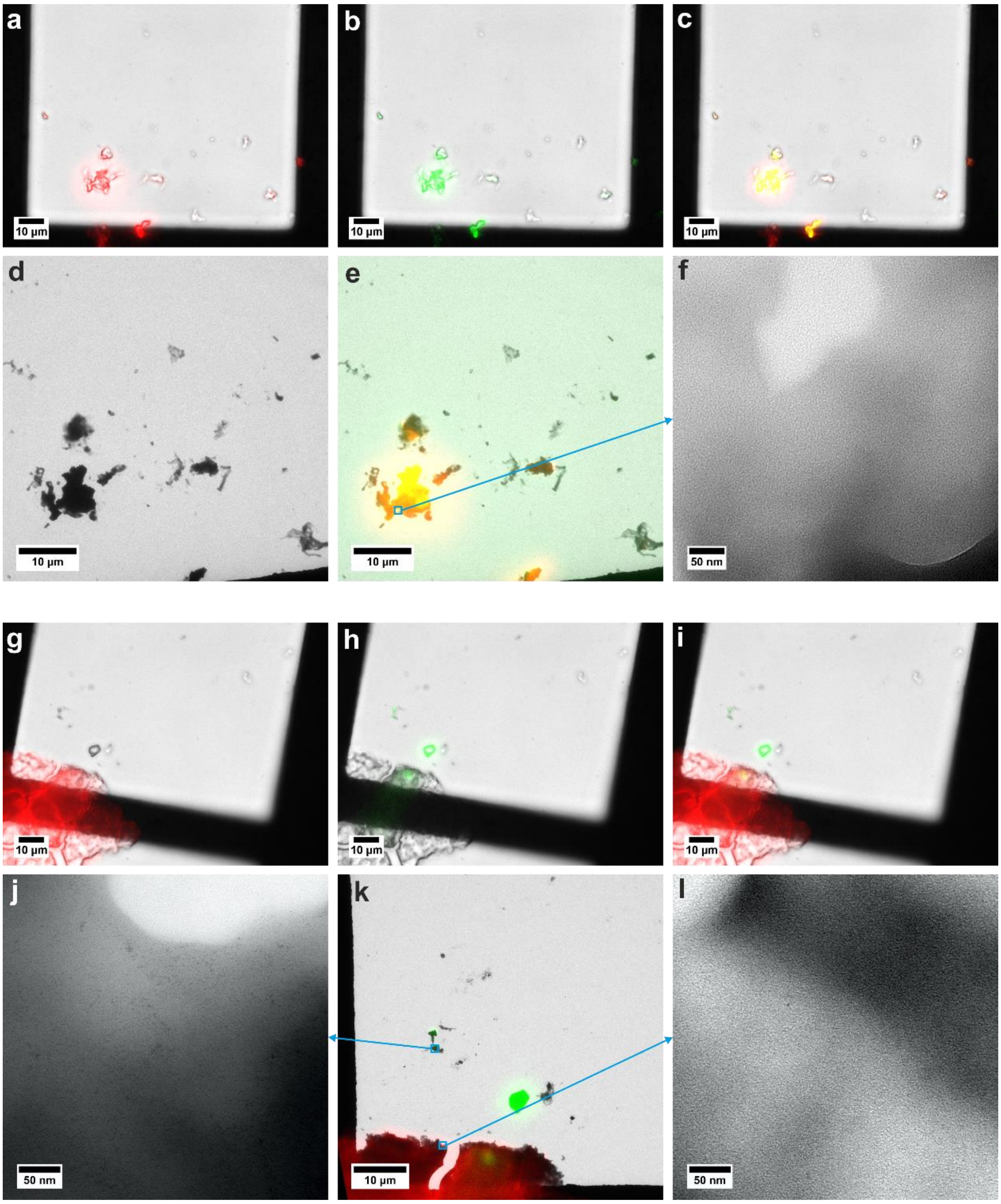

**Figure S6. Representative light and electron microscopy of fluorescently labelled peptide mixing**. 5(6)-carboxyfluorescein (cfl, green) and 5(6)-carboxytetramethylrhodamine (tmr, red) labelled distally pinned hairpins $_{pep}$cfl-HP-DT$_{dist}$ and $_{pep}$tmr-HP-DT$_{dist}$ respectively. These were mixed (a - f) when unfolded in 50% (v/v) acetonitrile and water or (h - l) 1 hour after rehydration in HBS. Optical microscope image overlaid with (a & g) red fluorescence, (b & h) green fluorescence, and (c & i) both channels. (d & j) A negatively stained TEM image, (e & k) negative stain TEM image overlaid with red and green fluorescence correlative light and electron microscopy (CLEM) image, and (f & l) negatively stained TEM image. Areas of higher magnification are indicated by cyan annotations. Fluorescently labelled peptides (10%) were mixed with non-labelled peptide $_{pep}$HP-DT$_{dist}$ (90%) in 50% (v/v) acetonitrile /water to make working stocks, which were freeze dried. For (a - f), the cfl and tmr working stocks were mixed when unfolded in 50% acetonitrile, and then freeze dried. The aliquots were hydrated in HBS (25 mM HEPES, 25 mM NaCl, pH 7.2) to a final concentration of 100 µM and incubated to assemble for 1 hour. For (h - l) the separate working stocks were hydrated in HBS to a final concentration of 100 µM and incubated to assemble for 1 hour, then mixed and incubated for 1 hour. 5 µL was applied to a TEM finder grid, washed, and dried, imaged on the fluorescence microscope, stained with 1% (w/v) aqueous uranyl acetate, washed, dried, and imaged in TEM. It was not possible to resolve FFTs of these assemblies, but a latticework pattern can be discerned in the images.

**Table S3. DNA primers used in this study**. Primers are paired to amplify sequences of interest to

| # | | sequence | $T_m$ (°) |
|---|---|---|---|
| Amplify sequences from GeneART delivery vectors | | | |
| 1 | F | 5' TCACTATAGGGCGAATTGGC 3' | 63.1 |
| 2 | R | 5' TTATTAACTAGTGTTTGAAGCTTATCAGTGG 3' | 63.3 |
| Introduce NcoI and SpeI restriction sites to GFP sequence Addgene vector | | | |
| 3 | F | 5' TAACCATGGGCATGAGCAAAGGCGAAGAGCTGTTC 3' | 59.1 |
| 4 | R | 5' TTATTACCATGGTTTGTACAGTTCATCCATACC 3' | 55.9 |

**Table S4. Cell strains used in this study**. Genes listed signify mutant alleles. Genes on the F' episome are wildtype unless indicated otherwise.

| **strain** | **genotype** |
|---|---|
| XL10-Gold® | Tet$^r$Δ(*mcrA*)*183* Δ(*mcrCB-hsdSMR-mrr*)*173 endA1 supE44 thi-1 recA1 gyrA96 relA1 lac Hte* [F´ *proAB lacI*$^q$*Z*Δ*M15* Tn*10* (Tet$^r$) Amy Cam$^r$] |
| SHuffle® T7 Express | *fhuA2 lacZ*::*T7 gene1 [lon] ompT ahpC gal λatt*::*pNEB3-r1-cDsbC* (*Spec*$^R$, *lacI*$^q$) Δ*trxB sulA11 R*(*mcr-73*::*miniTn10*—*Tet*$^S$)*2 [dcm] R(zgb-210::Tn10 --Tet*$^S$) *endA1* Δ*gor* Δ(*mcrC-mrr*)*114::IS10* |

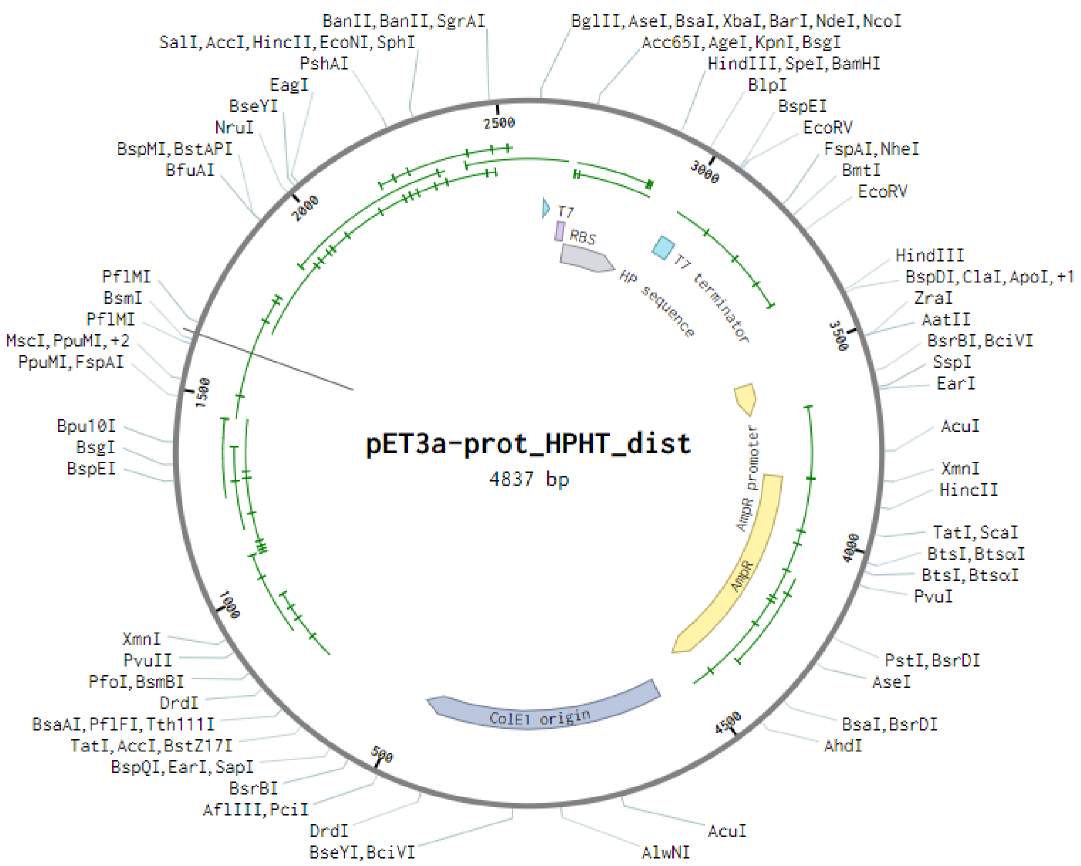

**Figure S7a. Plasmid vector map for pET3a vector containing hairpin sequence**. Labelled are: T7 promoter (T7), ribosome binding site (RBS), sequence for $_{pro}$HP-HT$_{dist}$, (HP sequence), T7 terminator (T7 terminator), ampicillin promoter (AmpR promoter), ampicillin resistance (AmpR), origin of replication (ColE1 origin).

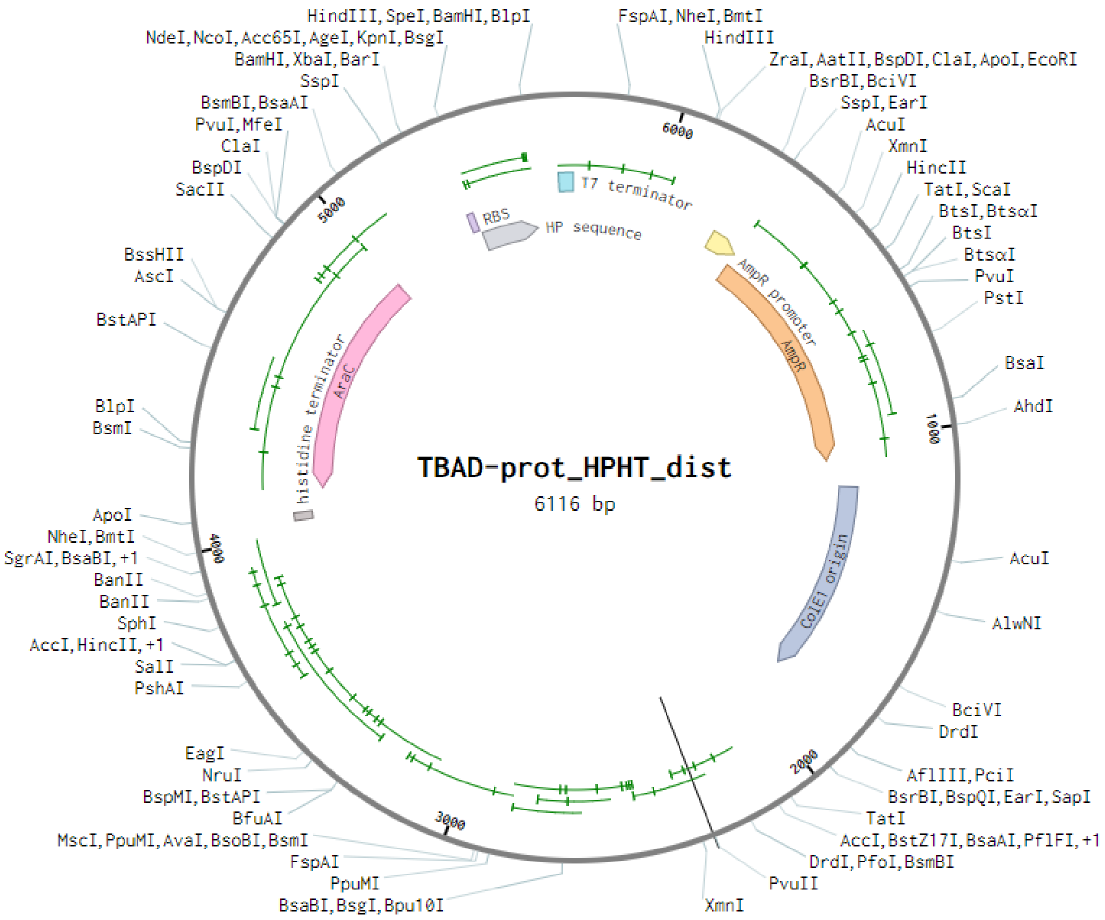

**Figure S7b. Plasmid vector map for TBAD vector containing hairpin sequence**. Labelled are: ribosome binding site (RBS), sequence for $_{pro}$HP-HT$_{dist}$, (HP sequence), T7 terminator (T7 terminator), ampicillin promoter (AmpR promoter), ampicillin resistance (AmpR), origin of replication (ColE1 origin), *AraC* for arabinose induction (AraC) and it's histidine terminator (histidine terminator).

**Table S5. DNA sequences of plasmids and inserts and corresponding protein sequence of inserts**. Homodimer sequence shown in grey, homotrimer sequence in green, cysteines for forming disulfide underlined.

| | **DNA and protein sequence** |
|---|---|
| $_{pro}$HP-$HT_{dist}$ | ATGTCAGGTTCCATGGGTTCTTCTGGTACCGGTGAAATTGCAGCAC-<br>TGAAATGTAAAAATGCAGCCCTGGAACAAGAGATCGCTGCACTGAAACAAGGTAGCGGTAG-<br>TGGCGAAATTGCCGCAATCAAAAAAGAAATCGCAGCCATCAAATGTGAGATCGCAG- |
| | **MSGSMGSSGT-<br>GEIAALKCKNAALEQEIAALKQGSGSGEIAAIKKEIAAIKCEIAAIKQGYYGSGHHHHHH** |
| $_{pro}$HP-$HT_{prox}$ | ATGTCAGGTTCCATGGGTTCTTCTGGTACCGGTGAAATTGCAGCAC-<br>TGAAACAAAAAAATGCAGCCCTGGAATGTGAGATCGCTGCACTGAAACAAGGTAGCGGTAG-<br>TGGCGAAATTGCCGCAATCAAATGTGAAATCGCAGCCATCAAAAAAGAGATCGCAG- |
| | **MSGSMGSSGT-<br>GEIAALKQKNAALECEIAALKQGSGSGEIAAIKCEIAAIKKEIAAIKQGYYGSGHHHHHH** |
| $_{pro}$GFP-HP-$HT_{dist}$ | ATGTCAGGTTCCATGGGCATGAGCAAAGGTGAAGAACTGTTTACCGGTGTTGTTCCGAT-<br>TCTGGTTGAACTGGATGGTGATGTTAATGGCCACAAATTTTCAG-<br>TTCGTGGTGAAGGCGAAGGTGATGCAACCAATGGTAAACTGACCCTGAAATTTATCTGTAC-<br>CACCGGCAAACTGCCGGTTCCGTGGCCGACACTGGTTACCACACTGACCTATGGTGTTCAG-<br>TGTTTTAGCCGTTATCCGGATCACATGAAACGCCACGATTTTTTCAAAAGCGCAATGCCG-<br>GAAGGTTATGTTCAAGAACGTACCATCTCCTTTAAAGATGATGGCAC-<br>CTATAAAACCCGTGCCGAAGTTAAATTTGAAGGTGATACCCTGGTGAATCGCATT-<br>GAACTGAAAGGCATCGATTTCAAAGAAGATGGTAATATTCTGGGCCACAAGCTGGAA-<br>TATAATTTCAATAGCCACAACGTGTATATCACCGCAGACAAACAGAAAAATGGCATCAAA-<br>GCCAACTTTAAGATTCGGCATAATGTTGAAGATGGCAGCGTTCAGCTGG-<br>CAGATCATTATCAGCAGAATACCCCGATTGGTGATGGTCCGGTTCTGCTGCCGGA-<br>TAATCATTATCTGAGCACCCAGAGCGTTCTGAGCAAAGATCCGAATGAAAAAC-<br>GTGATCACATGGTGCTGCTGGAATTTGTTACCGCAGCAGGTATTACACATGG- |
| | **MSGSMGMSKGEELFTGVVPILVELDGDVNGHKFSVRGEGEGDATNGKLTLKFICTT-<br>GKLPVPWPTLVTTLTYGVQCFSRYPDHMKRHDFFKSAMPEGYVQERTISFKDDGTYKTRAEV-<br>KFEGDTLVNRIELKGIDFKEDGNILGHKLEYNFNSHNVYITADKQKNGIKANFKIRHN-<br>VEDGSVQLADHYQQNTPIGDGPVLLPDNHYLSTQSVLSKDPNEKRDHMVLLEFVTAAGITHGMDELYKGT-<br>GEIAALKCKNAALEQEIAALKQGSGSGEIAAIKKEIAAIKCEIAAIKQGYYGSGHHHHHH** |

**Table S6. Gel compositions for SDS-PAGE**. Volumes of buffers and reagents required to make gels for SDS-PAGE analysis of proteins. Buffer compositions are listed in Table S7.

| **gel** | **component** | **stacking gel (mL)** | **separating gel (mL)** |
|---|---|---|---|
| tricine | water<br>50% (v/v) glycerol<br>gel buffer<br>40% 19:1 acrylamide/bis<br>APS<br>TEMED | 3.00<br>0.00<br>1.25<br>0.67 [5%]<br>0.06<br>0.01 | 0.62<br>2.60<br>3.33<br>3.33 [13%]<br>0.10<br>0.015 |
| glycine | water<br>gel buffer (1/2)<br>40% acrylamide/bis<br>10% (w/v) SDS<br>APS<br>TEMED | 3.00<br>1.25<br>0.67 [5%]<br>0.05<br>0.06<br>0.01 | 4.40<br>2.50<br>3.00 [12%]<br>0.10<br>0.10<br>0.015 |

**Table S7. Gel buffer compositions**. Protocols for making buffers used to make and run SDS-PAGE gels listed in Table S6.

| **solution** | **volume / pH** | **component** | **Mass / volume** |
|---|---|---|---|
| tricine gel buffer | 100 mL<br><br>pH 8.45 | 3.0 M Tris<br>SDS<br>(adjust with HCl) | 36.42 g<br>0.30 g |
| 1x cathode buffer | 500 mL<br><br>pH 8.25-8.30 | 0.10 M Tris<br>0.20 M tricine<br>0.10% SDS<br>(check pH correct) | 6.06 g<br>8.56 g<br>0.50 g |
| 1x anode buffer | 1000 mL<br>pH 8.9 | 0.2 M Tris<br>(adjust with HCl) | 24.22 g |
| glycine gel buffer 1 | 100 mL<br>pH 8.8 | 1.5 M Tris<br>(adjust with HCl) | 18.21 g |
| glycine gel buffer 2 | 100 mL<br>pH 6.8 | 0.5 M Tris<br>(adjust with HCl) | 6.07 g |
| 10x running buffer | 1000 mL<br><br><br>pH 8.3 | 0.25 M Tris<br>2.0 M glycine<br>SDS<br>(check pH correct) | 30.0 g<br>144.0 g<br>10.0 g |
| 4x gel loading buffer | 7.5 mL<br><br><br>If not blue add 1 drop 1 M NaOH | 0.4 M DTT<br>SDS<br>bromophenol blue<br>tricine gel buffer 2<br>glycerol<br>water | 0.463 g<br>0.600 g<br>0.030 g<br>3.00 mL<br>2.40 mL<br>up to 7.5 mL |
| Coomassie stain | 500 mL | Coomassie blue R250<br>water<br>ethanol<br>glacial acetic acid | 1.25 g<br>225 mL<br>225 mL<br>50 mL |
| destain | 1000 mL | water<br>glacial acetic acid<br>methanol | 700 mL<br>100 mL<br>200 mL |

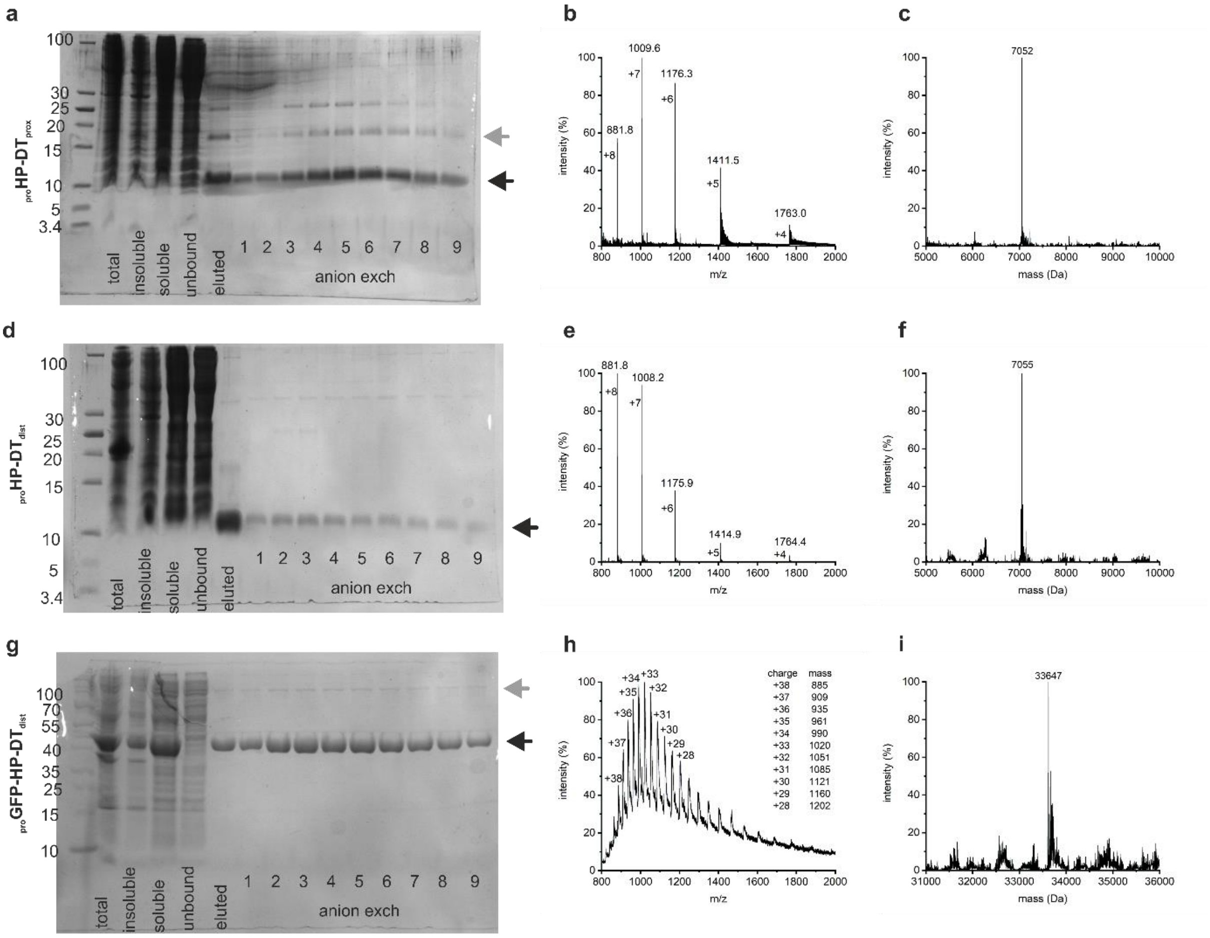

**Figure S8. Images of SDS-PAGE gels and nanospray mass spectra of purified protein**. (a, d & g) SDS page gels, (b, e & h) raw and (c, f & i) fitted nanospray mass spec for each expressed and purified protein. Each gel shows samples from the total cell lysate (total), the insoluble fraction (insoluble) and the soluble fraction (soluble), as well as the protein that did not bind to the Ni-NTA IMAC resin (unbound), that which was eluted by imidazole (eluted), and the fractions from anion exchange (anion exch). Tricine gels for low molecular weight proteins are shown in (a) $_{prot}$HP-HT$_{prox}$ and (d) $_{prot}$HP-HT$_{dist}$ (mass ~7 kDa, runs at ~10 kDa) with a low molecular weight ladder also labelled. Standard glycine gel is shown in (g) $_{prot}$GFP-HP-HT$_{dist}$.(mass ~40 kDa, runs at ~45 kDa). Monomer masses are indicated with a black arrow, dimer mass with a grey arrow. Proteins run slightly heavy on gels, probably because they are soluble and helical, which causes them to migrate as if they are slightly heavier when compared to the protein standard ladders.[17]

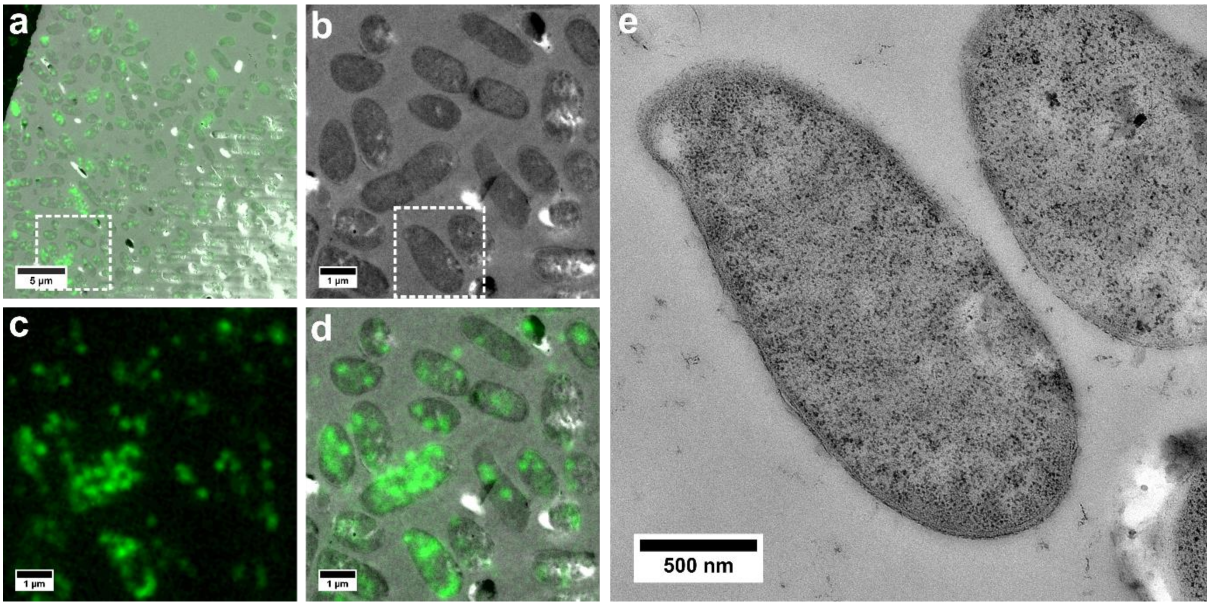

**Figure S9. Light and electron microscopy of immunolabelled cell sections**. High pressure frozen cells overexpressing $_{pro}$HP-DT$_{dist}$ under arabinose induction from the TBAD vector were sectioned, minimally stained and immunolabelled with Alexa fluor 488 (green) against the His-tag on the hairpin construct. Areas of higher magnification are indicated by dotted square. (a) Low magnification TEM with fluorescence overlay. Higher magnification separated out to show (b) TEM, (c) fluorescence and (d) CLEM overlay. (e) High magnification TEM image of area indicated in (b). Cells were sampled 6 hours after introduction to autoinduction media (LB + 0.1% arabinose, 0.1% glucose). 70 nm sections stained with anti-His antibody (1:500), then a secondary fluorescently labelled antibody (1 μg $mL^{-1}$) before imaging on the light microscope and then electron microscope.

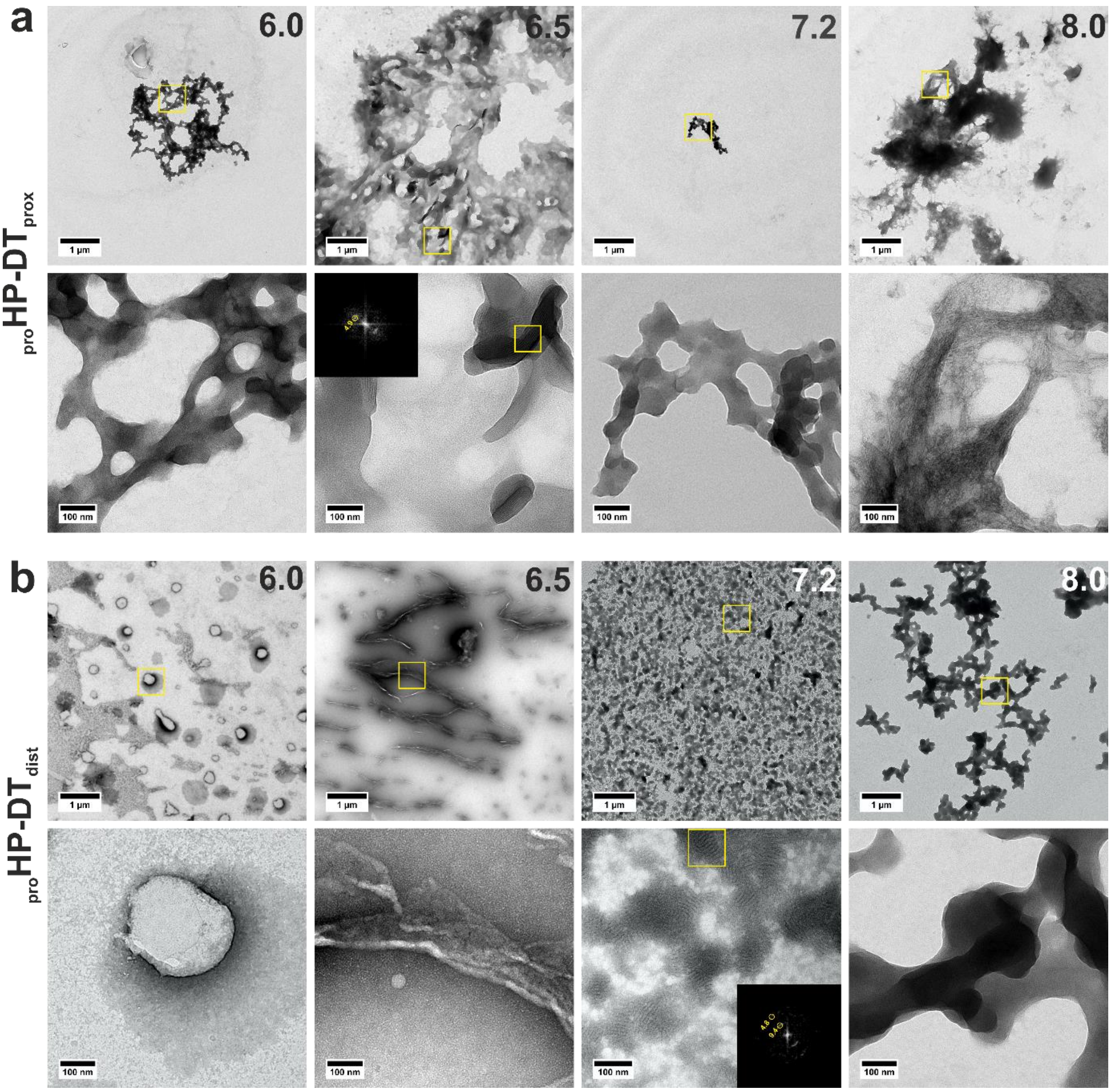

**Figure S10**. **TEM images of protein hairpins self-assembled at different pH**. (a) $_{pro}$HP-DT$_{prox}$ and (b) $_{pro}$HP-DT$_{dist}$ assembled in HBS (25 mM HEPES, 25 mM NaCl) adjusted to pH 6.0 (left), 6.5 (center left), 7.2 (center right, standard pH for all other experiments), and 8.0 (right).

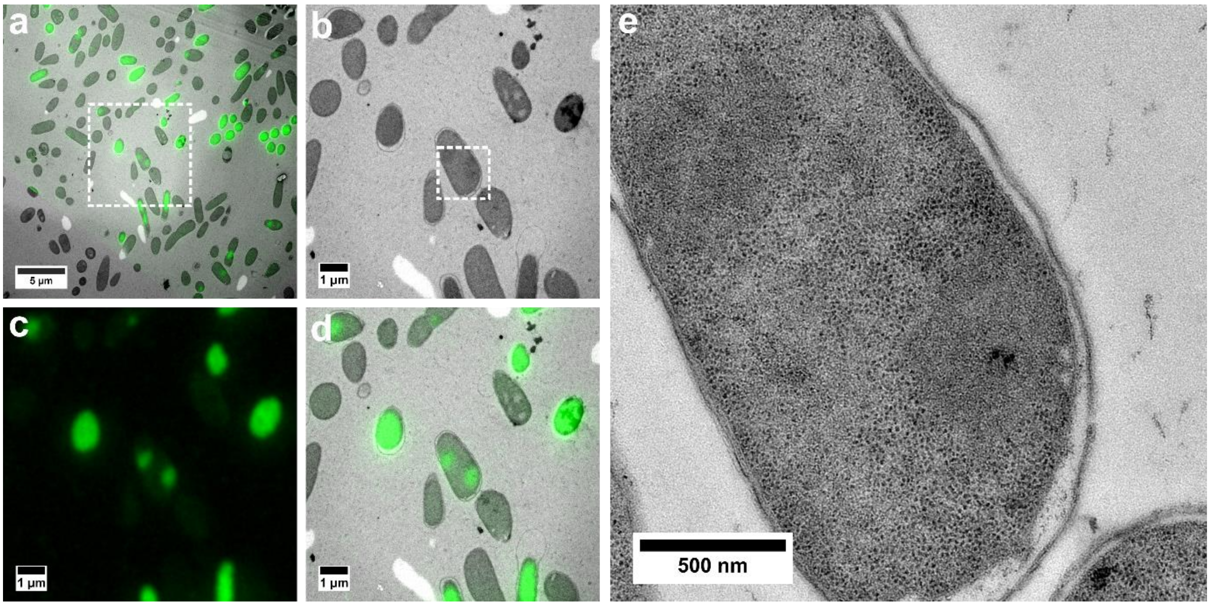

**Figure S11. Light and electron microscopy of cell sections expressing sfGFP – hairpin fusions**. High pressure frozen cells overexpressing $_{pro}$GFP-HP-DT$_{dist}$ under lactose autoinduction from the pET3a vector were sectioned, and minimally stained. Areas of higher magnification are indicated by dotted squares. (a) Low magnification TEM with fluorescence overlay. Higher magnification separated out to show (b) TEM, (c) fluorescence and (d) CLEM overlay. (e) High magnification TEM image of area indicated in (b). Cells were sampled 6 hours after introduction to autoinduction media (LB + 0.2% lactose, 0.05% glucose).

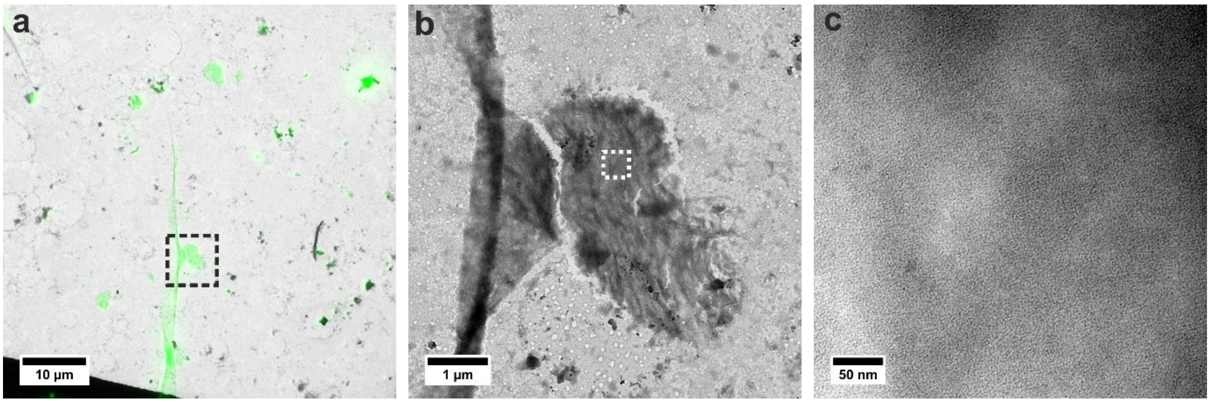

**Figure S12. CLEM of $_{pro}$GFP-HP-DT$_{dist}$ assembled *in vitro***. (a) TEM image overlaid with fluorescence from sfGFP to make CLEM, (b).medium magnification and (c) higher magnification image of area indicated with dotted lines shows a network pattern similar to that observed in the untagged protein hairpins. Unfortunately, the contrast in these images was not sufficient to analyze by FFT. The aliquots were hydrated in HBS (25 mM HEPES, 25 mM NaCl, pH 7.2) to a final concentration of 100 µM and incubated to assemble for 1 hour.
